# *Foxg1* bimodally tunes *L1*-mRNA and -DNA dynamics in the developing murine neocortex

**DOI:** 10.1101/2023.08.10.552831

**Authors:** Gabriele Liuzzi, Antonello Mallamaci

## Abstract

*Foxg1* masters telencephalic development via a pleiotropic control of its articulation. *L1* is a large retrotransposon family expressed within CNS and suggested to contribute to its genomic plasticity. Foxg1 represses gene transcription, and *L1* elements share putative Foxg1 binding motifs, suggesting the former might limit telencephalic expression (and activity) of the latter. We tested such prediction, in vivo as well as in engineered primary neural cultures, by loss- and gain-of-function approaches. We showed that *Foxg1*-dependent, transcriptional *L1* repression specifically occurs in neopallial neuronogenic progenitors and post-mitotic neurons, where it is supported by specific changes in the *L1* epigenetic landscape. Unexpectedly, we also found that Foxg1 physically interacts with *L1*-mRNA and positively impacts on neonatal neopallium *L1*-DNA content, antagonizing the retrotranscription-suppressing activity exerted by Mov10 and Ddx39a helicases. To our knowledge, *Foxg1* is the first CNS patterning gene acting as a bimodal retrotransposon modulator, limiting and promoting *L1* transcription and amplification, respectively, within a specific domain of the developing mouse brain.

## INTRODUCTION

Foxg1 encodes for an evolutionarily ancient transcription factor driving the development of the anterior brain [1]. It promotes the activation of subpallial [2] and neo-paleo-pallial [3] programs. It regulates pallial stem cells fate choice, promoting neuronogenesis at expenses of gliogenesis [4–6]. Moreover, it commits neocortical neurons to distinct laminar identities [1,7–9]. Next, it stimulates neuronal morphological maturation [4,10–12], and electrical activity [12,13], being in turn transiently upregulated by the latter [13,14]. Experimental *Foxg1* knock-down in vivo reduces social interaction and results in selective impairment of specific learning and memory abilities [11,15,16]. In humans, several *FOXG1* copy number variations (CNVs) and structural mutations have been described. Heterozygosity for them results in a complex array of severe neuropathological scenarios, collectively referred to as FOXG1 syndrome, for which no cure is so far available [17–23]. Classically known as a transcriptional transrepressor [24], Foxg1 has been more recently implicated in straight control of extra-transcriptional functions, such as post-transcriptional ncRNA processing [25], translation [26] and mitochondrial biology [27].

Albeit tightly controlled [28,29], transposable elements including *L1s* are actively transcribed. In particular, specific ensembles of such elements are activated concomitantly with distinct, early histogenetic routines [30], and, in some cases, their transcription is needed for the progression of these routines [31–33]. Moreover, a subset of full-length *L1s* is able to undergo somatic retrotransposition. As little as about 100 in humans, such retrotransposition-competent *L1s* are about 3000 in mice, 900, 400 and 1800, belonging to A, Gf and Tf sub-families, respectively [34,35], and products of their somatic retro-transposition generally miss family-specific 5’UTRs [36], while retaining shared orf2 and 3’UTR regions. Clonal analysis robustly demonstrated that somatic retrotransposition takes place within the developing embryo at different times and in variety of cell types, with special emphasis on the developing CNS [30,37–40]. The magnitude of L1 neo-retrotransposition in the human CNS has been hotly debated. In particular, human neocortical/ hippocampal neurons have been reported to harbor somatic L1 insertions at a frequencies estimated between 0.2 and 80 events per neural cell [41–45]. In the mouse an increase of *L1*-DNA content has been reported to occur as well, in both neocortical and hippocampal neurons from E15.5 and P14, close to +30% [46]. A substantial fraction of Foxg1 protein is stably bound to chromatin [47], suggesting it might be implicated in long term gene repression. Next, motif enrichment analysis (MEA) by Jaspar software [48] revealed a high score, putative Foxg1 binding site (RTAAACAW) within *L1*-*orf2* cds (our unpublished data). Based on that, we hypothesized that Foxg1 may be implicated in control of *L1* transcription. We confirmed this prediction, modeling its articulation within the embryonic neocortex, along the neuronogenic lineage. We showed that Foxg1-dependent L1 repression mainly occurs in neuronogenic progenitors and post-mitotic neurons, where it is supported by specific changes in the epigenetic landscape. Unexpectedly, we also found that Foxg1 positively impacts on neopallial *L1*-DNA content, antagonizing the retrotranscription-suppressing activity exerted by Mov10 and Ddx39a helicases.

## RESULTS

### In vivo *Foxg1* down-regulation of *L1*-mRNA

To assess if Foxg1 is implicated in control of *L1* trascription, we compared *L1*-mRNA levels in the neocortex of P0 *Foxg1^−/+^* mice [49] with wild type controls. As expected, we found a substantial *L1* upregulation in these mutants. It characterized the diagnostic "*L1*.orf2" amplicon, shared by all *L1* sub-families (+19.7±1.6%, p<0.014, n=7,7), as well "*L1*.5’UTR.A, .Gf, and .Tf" amplicons, peculiar to the three corresponding transposition-competent sub-families (+17.2±0.3%, p<0.008, n=7,7; +33.0±0.1%; p<10^−4^, n=7,7; +15.8±0.5%, p<0.006, n=7,7; respectively) (**Fig. 1**).

**Figure 1.**
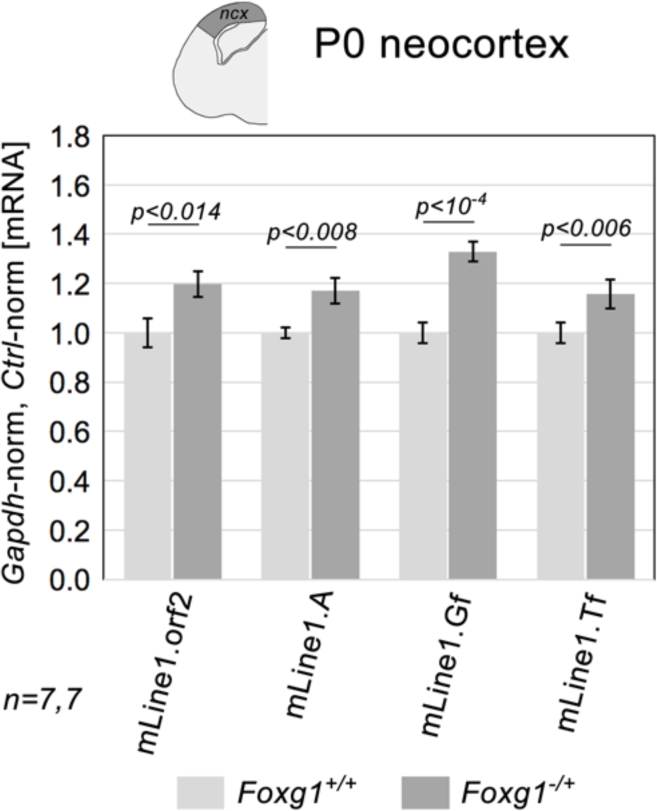
L1 transcripts levels in neocortex of *Foxg1^−/+^* mouse neonates and controls. RT-PCR quantitation of pan-*L1* diagnostic amplicon "orf2", and family-specific amplicons, "A", "Gf" and "Tf". Data double normalized, against *Gapdh* and wild-type controls. Error-bars representing sem’s. Statistical significance of results evaluated by t-test (one-tailed, unpaired). *n* is the number of biological replicates, i.e. neocortices taken from distinct pups.

### In vitro modeling of *L1-mRNA* progression in murine developing neocortex

To ease dissection of Foxg1 control of *L1* expression, we developed three protocols, "type-I", "-II" and "-III", to generate primary neural cultures representative of early, mid and late phases of pallial neuronogenesis, respectively (**Fig. 2A**). Displaying progressively longer durations, such protocols differed for terminal exposure of neural cells to "pure pro-proliferative", "mixed pro-proliferative/pro-differentiative", and "pure pro-differentiative" media, respectively. Type I cultures gave substantial fractions of Sox2^+^Tubb3^−^ presumptive NSCs (39.3±0.9%; n=3) and Sox2^−^Tubb3^−^ neuronogenic progenitors (30.1±0.7%; n=3), with limited Tubb3^+^ neuronal output (30.6±1.2%; n=3). Prevalence of these two precursors was lowered (to 22.2±1.4% and 16.6±1.1%, respectively; n=3) in Type II cultures, characterized by more frequent neurons (61.2±2.0%; n=3). As expected, neuronal prevalence further increased in Type III cultures (71.7±1.0%; n=3).

**Figure 2.**
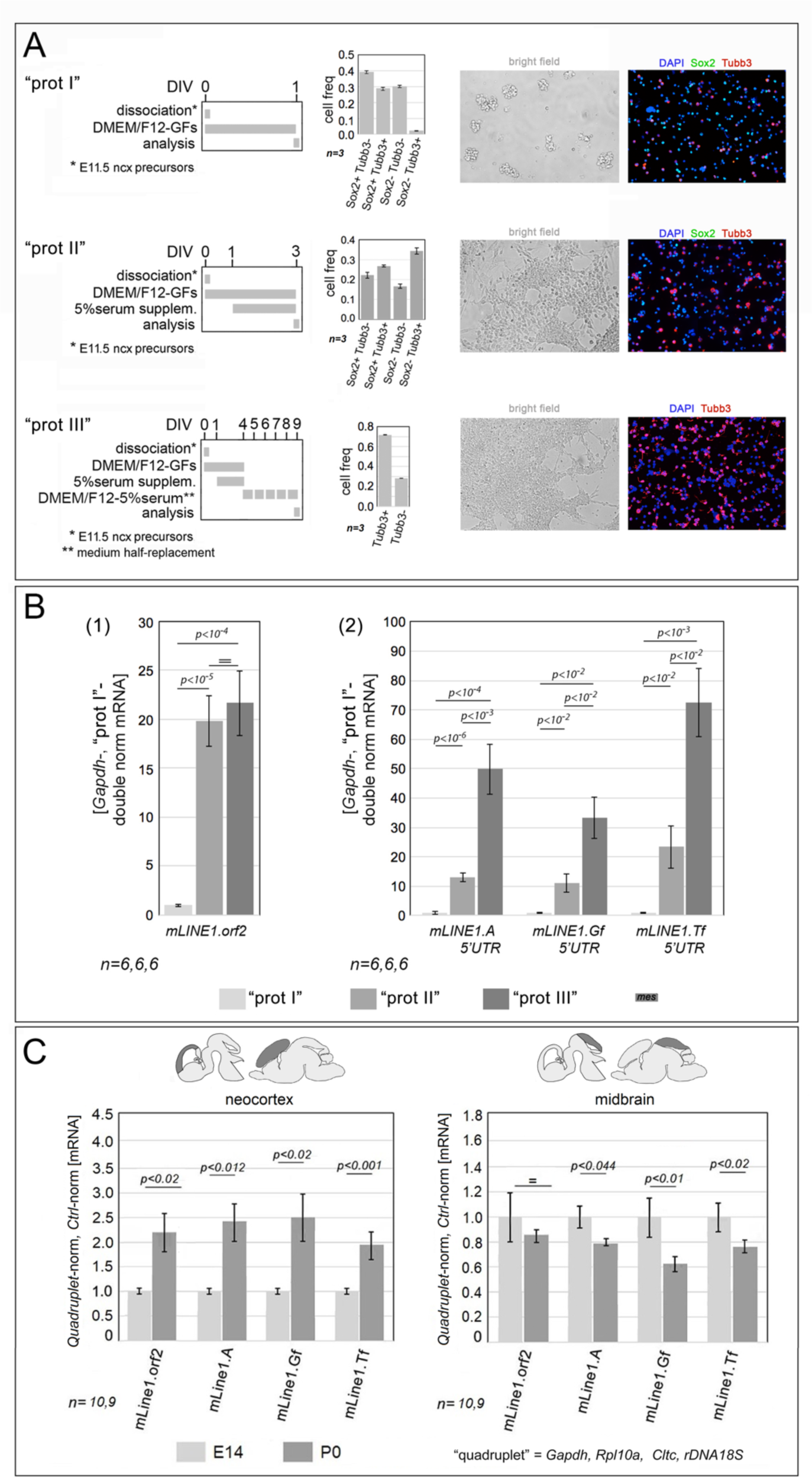
In vitro modeling of *L1*-mRNA progression in murine developing neocortex. In (**A**) shown are three protocols aimed at generating primary cultures representing early, mid and late phases of neuronogenesis. In such cultures, neural cells are terminally exposed to GFs, GFs and serum, and serum, respectively. Hence, they: (I) are enriched in neural stem cells (NSCs, Sox2^+^Tubb3^−^) and neuronogenic progenitors (NPs, Sox2^−^Tubb3^−^), (II) include comparable fractions of NSCs, NPs and neurons (Ns, Tubb3^+^), and (III) are highly enriched in Ns, respectively. In (**B**) provided are RT-PCR quantitations of pan-*L1* diagnostic amplicon "orf2" (1), and family-specific amplicons, "A", "Gf" and "Tf" (2), in neural cultures set according to protocols I, II and III. Data double normalized, against *Gapdh* and "protocol I" values. In (**C**) shown are RT-PCR quantitations of the same diagnostic amplicons in neocortex and mesencephalic tectum at embryonic day 14.5 (E14.5) and birth (P0). Data double normalized against the geometric mean of *Gapdh*, *Rpl10a*, *Cltc* and *rDNA18S* ("quadruplet") and "E14.5" values. In (B,C) error-bars representing sem’s. Statistical significance of results evaluated by t-test (one-tailed, unpaired). *n* is the number of biological replicates, i.e. independently cultured cell aliquots, originating from pooled, wild-type E11.5 neocortical primordia (B) or distinct embryos/pups (C).

Next, we profiled these cultures for *L1*-mRNA expression levels, by qRT-PCR (**Fig. 2B**). An increasing progression, from "type I" up to "type III" ones, emerged at the pan-*L1* diagnostic amplicon "*L1.orf2*" (specifically, "type I" culture-normalized values were: 1.00±0.11, 19.85±2.59, and 21.65±3.27, respectively, with *p*_(I-vs-II)_<10^−5^, *p*_(II-vs-III)_<0.34, *n*=6,6,6). Similar progressions were also detectable at sub-family-specific amplicons, "*L1.5’UTR.A* ("type I" culture-normalized values: 1.00±0.39, 13.01±1.49, and 49.87±8.57, respectively, with *p*_(I-vs-II)_<10^−5^, *p*_(II-vs-III)_< 10^−3^, *n*=6,6,6), *.Gf* ("type I" culture-normalized values: 1.00±0.10, 11.16±3.10, and 33.31±7.14, respectively, with *p*_(I-vs-II)_<10^−2^, *p*_(II-vs-III)_< 10^−2^, *n*=5,6,6), and *.Tf*" ("type I" culture-normalized values: 1.00±0.05, 23.41±7.26, and 72.48±11.49, respectively, with *p*_(I-vs-II)_<10^−2^, *p*_(II-vs-III)_< 10^−2^, *n*=5,6,5). All this points to a generalized *L1* upregulation, associated with progression of neocortical neuronogenesis.

To assess biological plausibility of these results, we repeated this analysis *in vivo*, comparing *L1* expression in neocortical tissue taken from E14.5 (mid-neuronogenic) and P0 (post-neuronogenic) mice (**Fig. 2C**). [Here, to strengthen the quality of results, we normalized *L1* qRT-PCR values against a specific "gene quadruplet". This included three RNA-pol II-transcribed genes (*Gapdh*, *Rpl10a* and *Cltc*), characterized by comparable expression profiles in apical precursors (APs), basal progenitors (BPs), early neurons (eNS) and late neurons (lNs) [50] (**Table S2A**), as well as by poor sensitivity to *Foxg1* manipulation [51] (**Table S2B**). It also included RNA-pol I*-*transcribed *rDNA-45S*, from which the large majority of cell RNA complement is generated]. As expected, *L1-mRNAs* were robustly upregulated in P0 compared to E14.5 neocortices (1), as detected by amplicons "*L1.orf2*" (+122.3±47.7%, p<0.004, n=10,7), "*L1.5’UTR.A*" (+140.6±44.0%, p<0.001, n=10,7), "*L1.5’UTR.Gf*" (+149.6±51.4%, p<0.003, n=9,7), and "*L1.5’UTR.Tf*" (+91.0±28.1%, p<0.001, n=10,7). As a specificity control, a similar analysis was performed on mesencephalic tectum harvested from the same animals. Intriguingly, this displayed an opposite E14.5 →P0 *L1*-mRNA dynamics (2): "*L1.orf2*": −15.0±8.2%, p<0.221, n=9,8; "*L1.5’UTR.A*": −20.9±5.1%, p<0.045, n=9,8; "*L1.5’UTR.Gf*": −36.5±4.7%, p<0.013, n=10,8; and "*L1.5’UTR.Tf*" −24.3±5.3%, p<0.051, n=10,8.

### Modeling *Foxg1* regulation of *L1*-mRNA

To dissect Foxg1 control of *L1* transcription, firstly, we evaluated *L1*-mRNA levels in "type II", mid-neuronogenic cultures, made constitutively Foxg1-LOF (see **Table S1**), by CRISPR-Cas9 technology and lentiviral transgenesis (**Fig. 3A**). Consistently with P0 *Foxg1^−/+^* pups (**Fig. 1**), neocortical *Foxg1*-LOF cultures displayed a systematic upregulation of *L1*-mRNAs, +25.9±7.4% (*p*<0.004), +21.6±10.9% (*p*<0.054), and +18.7±6.7% (*p*< 0.017), as for sub-families A, Gf and Tf, respectively, with *n*=10,11 (**Fig. 3A, S1A**).

**Figure 3.**
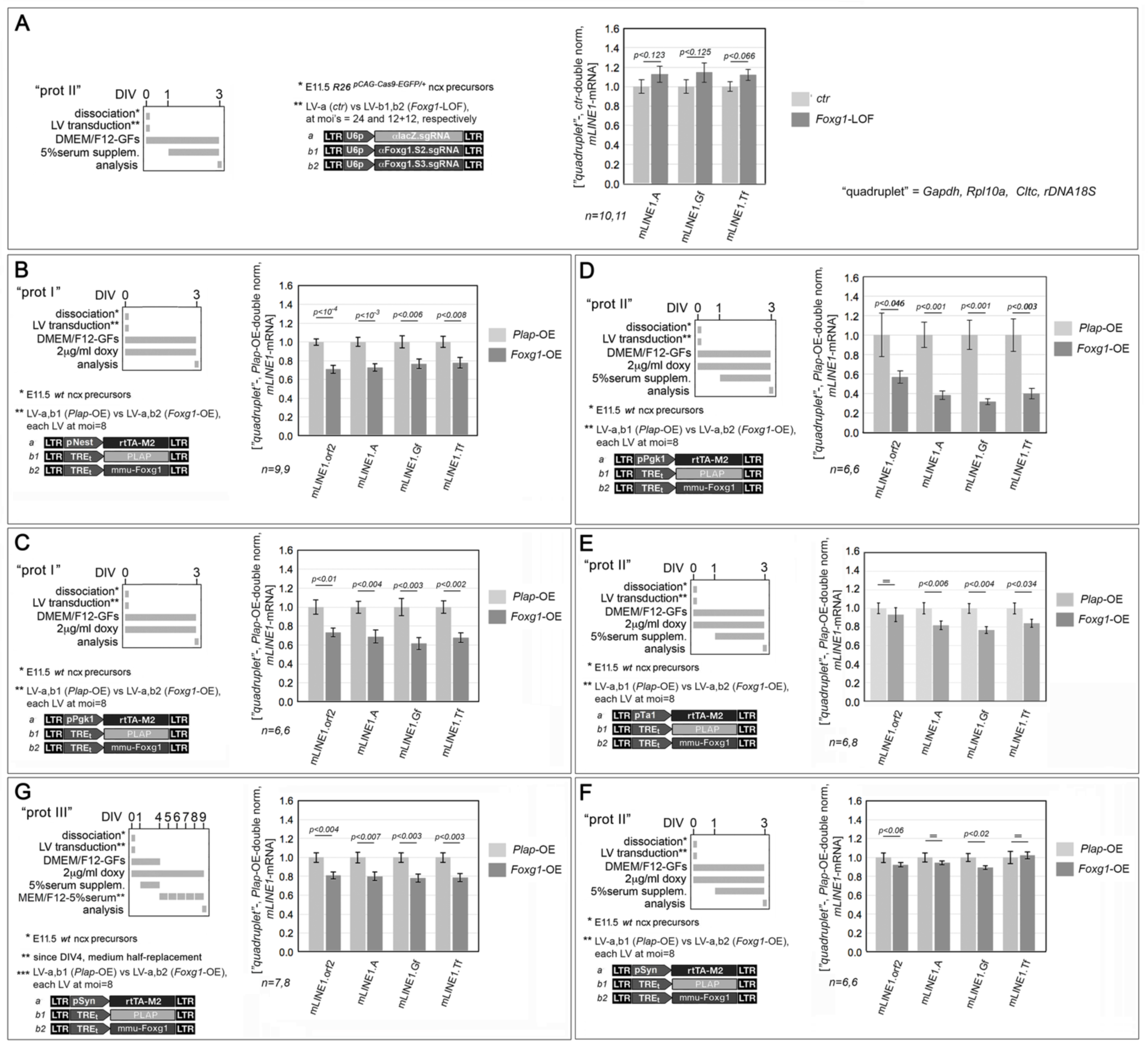
Impact of *Foxg1* manipulation on *L1*-mRNA levels, in progressively more advanced neuronogenic pallial cultures. In (**A)** and (**B-G**), shown are outcomes of *Foxg1* downregulation (*Foxg1*-LOF) and overexpression (*Foxg1*-OE), respectively, in early-(B,C), mid-(A,D-F) and late-(G) neuronogenic cultures, set according to type I, II and III protocols, respectively. To left, protocols and lentiviral vectors employed. Transgenes driven by pNes, pTα1 and pSyn promoters, active in NSCs, NPs/Ns and Ns, respectively, and ubiquitously firing U6p and pPgk1 promoters. To right, results. RT-PCR quantitation of pan-*L1* diagnostic amplicon "orf2", and family-specific amplicons, "A", "Gf" and "Tf", in neural cultures set according to above-mentioned protocols. Data double normalized against gene quadruplet (*Gapdh*, *Rpl10a*, *Cltc* and *rDNA 18S*) and control values. Error-bars representing sem’s. Statistical significance of results evaluated by t-test (one-tailed, unpaired). *n* is the number of biological replicates, i.e. independently cultured and engineered cell aliquots, originating from pooled *R26^pCAG-Cas9-EGFP/+^* (A) and *wild-type* (B-G), E11.5 neocortical primordia.

Next, we investigated *L1*-mRNA response to Foxg1 upregulation. To get insights into temporal and intra-neuronogenic lineage articulation of Foxg1 modulation of *L1* expression, we run multiple *Foxg1*-OE assays, employing different, early-mid- and late-neuronogenic, cultures, and restricting *Foxg1* transgene expression by means of ubiquitous pPgk1 promoter and cell-type specific, pNes, pTa1 and pSyn promoters, active in NSCs, NPs/Ns, and Ns, respectively [4,13].

Foxg1 overexression in early-neuronogenic ("type I") cultures, driven by either pNes or pPgk1, gave rise to a similar and generalized down-regulation of *mLINE*-mRNAs. Specifically, in case of pNes-manipulated cultures, L1.orf2, .A, .Gf, and .Tf signals were decreased, by 29.1±4.1% (*p*<10^−4^, *n*=9,9), 27.3±3.8% (*p*<10^−3^, *n*=9,9), 23.2±5.0% (*p*<0.006, *n*=9,9), and 22.3±5.5% (*p*<0.008, *n*=9,9), respectively (**Fig. 3B**), in case of pPgk1-manipulated ones, by 26.8±4.5% (*p*<0.006, *n*=6,6), 31.0±6.7% (*p*<0.004, *n*=6,6), 38.5±6.3% (*p*<0.003, *n*=6,6), and 32.2±5.0% (*p*<0.002, *n*=6,6), respectively (**Fig. 3C**).

Next, mid-neuronogenic ("type II") *Foxg1*-OE cultures also gave different results depending on the promoter driving the *Foxg1* transgene. The most prominent *L1* downregulation was observed in *Foxg1*-OE^pPgk1^ cultures. Here *L1*.orf2, .A, .Gf, and .Tf signals were decreased by 42.9±6.1% (*p*<0.046, *n*=6,6), 61.4±4.4% (*p*<0.001, *n*=6,6), 68.1±2.9% (*p*<0.001, *n*=6,6), 60.2±5.4% (*p*<0.003, *n*=6,6) (**Fig. 3D**). A milder decline of *L1-*mRNA was detectable in *Foxg1*-OE^pTa1^ cultures, where L1.orf2, .A, .Gf, and .Tf signals were decresed by 6.9±7.5% (*p*<0.238, *n*=7,5), 18.1±4.6% (*p*<0.025, *n*=7,5), 23.4±3.5% (*p*<0.004, *n*=7,5), and 16.1±4.3% (*p*<0.034, *n*=7,5), in Foxg1-OE^pTa1^ cultures (**Fig. 3E**). *L1*-mRNA levels were prevalently unaffected in *Foxg1*-OE^pSyn^ cultures, except *L1*.orf2 and .Gf signals, reduced by 9.6±0.4% (*p*<0.063, *n*=6,5), and 10.7±0.8% (*p*<0.024, *n*=6,5), respectively (**Fig. 3F**).

Last, late-neuronogenic cultures over-expressing *Foxg1* under the control of pSyn displayed a robust *L1*-mRNA down-regulation, with L1.orf2, .A, .Gf, and .Tf signals decresed by 19.2±3.8% (*p*<0.004, *n*=7,8), 14.3±6.7% (*p*<0.065, *n*=7,8), 15.6±7.2% (*p*<0.054, *n*=7,8), and 21.3±4.0% (*p*<0.003, *n*=7,7), respectively (**Fig. 3G**).

While pointing to a generalized *Foxg1*-dependent *L1* down-regulation, these results offer valuable hints about temporal and cell-type specific articulation of this process. In this respect, we have to distinguish between *late* neuronogenic cultures manipulated by a *pSyn-driven Foxg1* transgene (**Fig. 3G**) and *other cultures* (**Fig. 3B-F**). In the former case the promoter was active in cells occupying a terminal position along the neuronogenic sequence and the culture was allowed to age for a sufficiently long time, so that Foxg1 protein robustly arose within the same cell type where the promoter fires. Conversely, in the other cases, the promoters were active in transient precursor types, within relatively short-lived cultures. In this way, Foxg1 protein arousal could have taken place in a cell type where the promoter is no longer active (or there might have been not enough time to get a pronounced protein upregulation).

In the light of these considerations, the interpretation of data obtained in Foxg1-OE^pSyn^, late neuronogenic cultures is straightforward, pointing to a consistent *neuronal* inhibition of *L1* elements belonging to all sub-families by Foxg1. This inference was corroborated by the results of supplemental *Foxg1* manipulations, upwards and downwards, constitutively performed in pure neuronal cultures fully depleted of glial cells by araC supplementation (**Fig. 4**). In such neocortical neurons, *Foxg1* overexpression reduced the *L1*.orf2 qRT-PCR signal, by 25.2±8.7% (*p*<0.046, *n*=4,4) upon *Gapdh*-normalization (**Fig. 4**, graph (1)), by 33.5±8.7% (*p*<0.025, *n*=4,4) upon *Rpl10a*-normalization (**Fig. 4**, graph (2)), whereas *Foxg1*-knockdown increased such signal, by 66.4±21.9% (*p*<0.016, *n*=4,4) upon *Rpl10a*-normalization (**Fig. 4**, graph (3). Altogether, these results point to an overt key role played by Foxg1 in *physiological tuning* of neuronal *L1* transcription.

**Figure 4.**
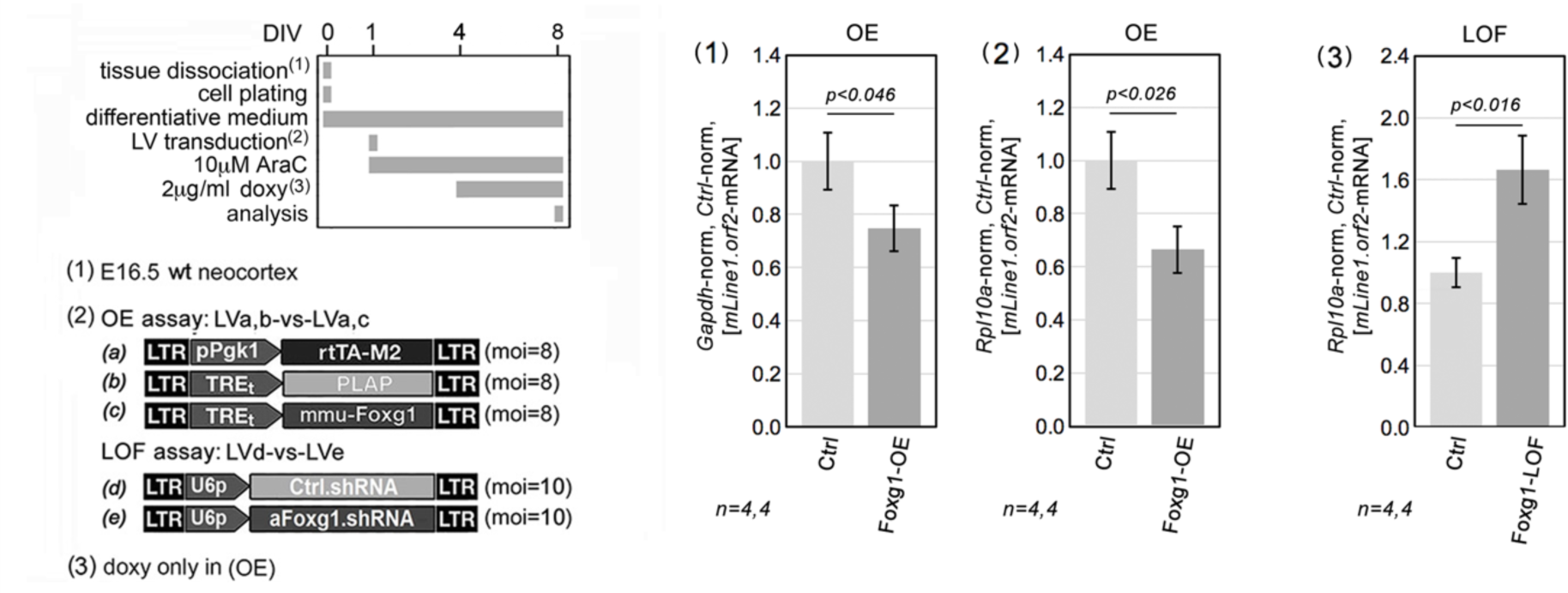
Impact of constitutive *Foxg1* overexpression on *L1*-mRNA levels, in primary, neuron-enriched cultures. To left, protocols and lentiviral vectors employed, to right, results. Neuronal enrichment obtaind by early araC supplementation. Foxg1 constitutively overexpressed (OE) by a pPgk1-driven transgene, or down-regulated (LOF) by RNAi. *L1*-mRNA data double normalized against *Rpl10a* (or *Gapdh*) and control samples. Scalebars, sem’s. Statistical evaluation of results by t-test (one-tail, unpaired). *n* = number of biological replicates, i.e. independently cultured and engineered cell aliquots originating from a common neural precursor pool.

Conversely, the phenotype of early- and mid-neuronogenic cultures overexpressing *Foxg1* required further disambiguation. To this aim, (1) we quantified the sizes of NSCs, NPs and Ns compartments of differently engineered cultures, (2) we scored Foxg1 protein levels within each compartment, and (3) we finally compared results of this analysis with *L1-*mRNA dynamics peculiar to the corresponding cultures.

We found that, *within early-neuronogenic preparations*, pNes-driven *Foxg1* elicited a prominent increase of NPs (f_NP_(*Foxg1*-GOF)=39.16±1.14% vs f_NP_(ctrl)=11.93±2.58%, p<10^−5^, n=6,5) at expenses of NSCs (f_NSC_(*Foxg1*-GOF)=57.81±1.53% vs f_NSC_(ctrl)=84.58±2.81%, p<10^−5^, n=6,5), while keeping Ns to a minimum (**Fig. 5A** (1)). Moreover, within both NSCs and NPs compartments, it specifically increased the frequency of precursors expressing Foxg1 at the highest levels (i.e. belonging to the first decile), with ^NSC^f_dec1_(ctrl)=3.4% and ^NSC^f_dec1_(*Foxg1*-GOF)=15.4%, and ^NP^f_dec1_(ctrl)=48.5% and ^NP^f_dec1_(*Foxg1*-GOF)= 83.3%, respectively (**Fig. 5A** (2)). In this way, among additional cells falling within the first-decile upon *Foxg1* overexpression, >4/5 belonged to the NP compartment (Δf_NP*dec1_ = f_NP_(*Foxg1*-GOF) * ^NP^f_dec1_(*Foxg1*-GOF) - f_NP_(ctrl) * ^NP^f_dec1_(ctrl) = 0.392 * 0.832 - 0.119 * 0.485 = 0.268), and less than 1/5 to the NSC one (Δf_NSC*dec1_ = f_NSC_(*Foxg1*-GOF) * ^NSC^f_dec1_(*Foxg1*-GOF) - f_NSC_(ctrl) * ^NSC^f_dec1_(ctrl) = 0.578 *0.154 - 0.846 * 0.034 = 0.060). In other words, the most prominent Foxg1 overexpression elicited upon delivery of a pNes-driven *Foxg1* transgene to early-neuronogenic cultures took mainly place in NPs. That suggests that, within these cultures, *L1*-mRNA downregulation evoked by *^pNes^Foxg1*-OE (**Fig. 3B**) likely occurred just in such progenitors, and that NSCs contribution to this phenomenon was marginal, if any (**Fig. 7** (1)).

**Figure 5.**
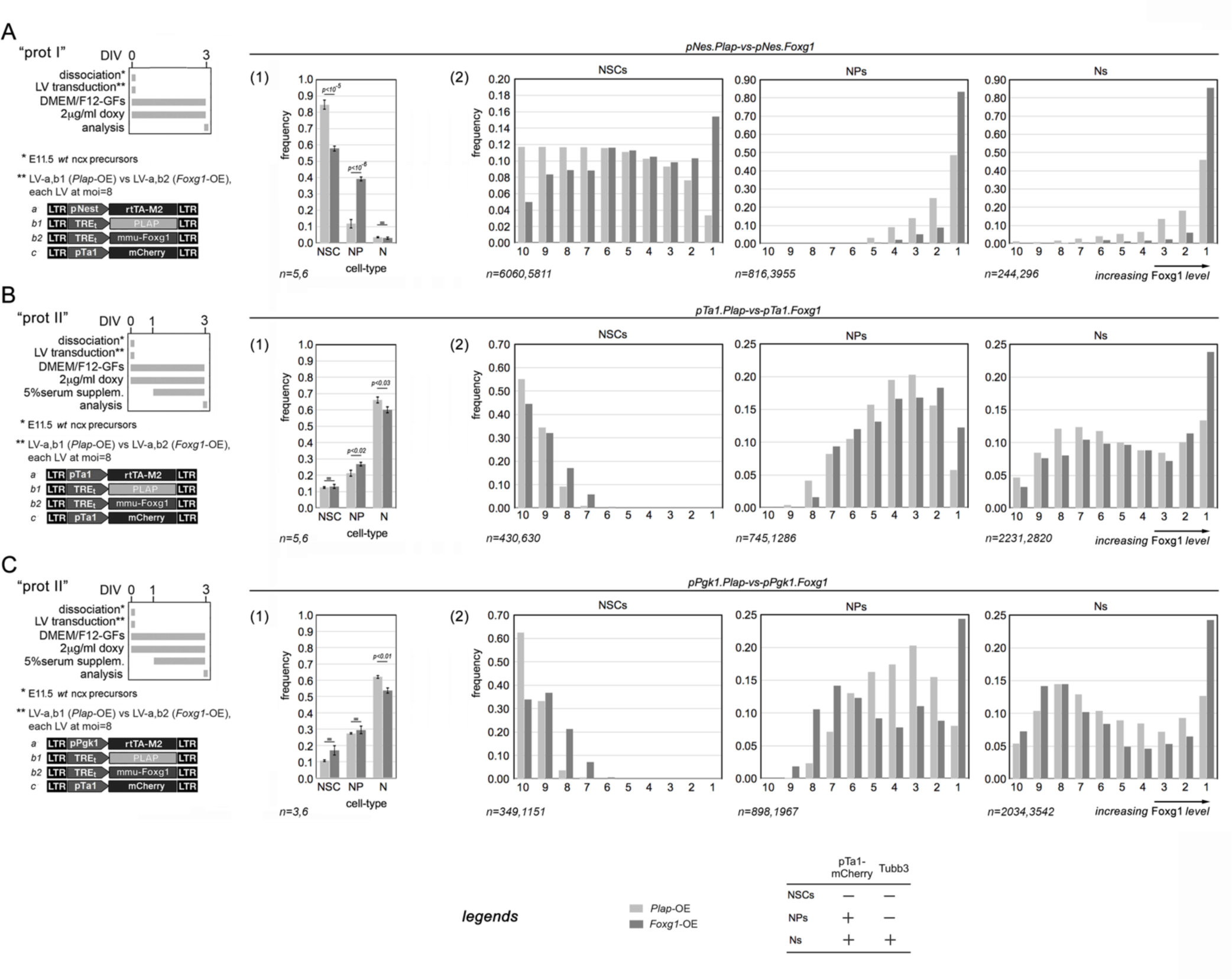
Quantification of Foxg1 protein levels in distinctive neural precursor types, upon *Foxg1* over-expression driven by lentiviral vectors and cell-type specific promoters. Throughout the figure, to left, protocols and lentiviruses employed, to right, results. Assays run in early- (**A**), mid-(**B,C**) and late-(**D**) neuronogenic cultures, set according to type I, II and III protocols, respectively. Cultures over-expressing *Foxg1* or a control (*Plap*). Foxg1-encoding transgenes driven by pNes (A) and pTa1 (B) promoters, active in NSCs, and NPs/Ns, respectively, and ubiquitously firing pPgk1 promoter (C). NSCs, NPs and Ns recognized on the basis of their ^pTa1^mCherry^−^/Tubb3^−^, ^pTa1^mCherry^+^/Tubb3^−^, and ^pTa1^mCherry^±^/Tubb3^+^ profiles, respectively. Foxg1 cell content evaluated by quantitative immuno-fluorescence. In graphs (1), shown are prevalences of NSCs, NPs and Ns in neural cultures set according to the different protocols. Error-bars representing sem’s. Statistical significance of results evaluated by t-test (one-tailed, unpaired). *n* is the number of biological replicates, i.e. independently cultured and engineered cell aliquots, originating from pooled, *wild-type* E11.5 neocortical primordia. In graphs (2), shown are frequencies of NSCs, NPs and Ns falling within distinct Foxg1 expression deciles, i.e. normally equi-numerous bins characterized by decreasing (from 1 to 10) Foxg1 expression levels. n is the number of cells profiled, evenly pooled from the biological replicates referred to as for graph (1) series.

Next, we performed a similar analysis of *mid-neuronogenic preparations* harboring the pTα1- and pPgk1-driven *Foxg1* transgenes, i.e. those eliciting the strongest impact on *L1*-mRNA dynamics (**Fig. 3D-F**). We found that the two transgenes altered only marginally the sizes of the main culture compartments, both eliciting a moderate shrinkage of the neuronal one (from 66.1±1.9% to 60.0±1.8%, with *p*<0.03 and *n*=5,6, as well as 62.0±0.9% to 53.6±1.6%, with *p*<0.01 and *n*=3,6, respectively) (**Fig. 5B** (1) and **5C** (1)). More intriguingly, while similarly perturbing neuronal Foxg1 expression levels, they led to a different mis-distribution of Foxg1 protein in NPs. Specifically, upon *Foxg1*-OE, the increase of the NP aliquot falling in the first expression decile was much more pronounced in *^pPgk1^Foxg1*-OE cultures (0.244 - 0.080 = 0.164) compared to *^pTa1^Foxg1*-OE ones (0.122 - 0.058 = 0.064) (**Fig. 5B** (2) and **5C** (2)). Together with stronger *L1* inhibition elicited by pPgk1-driven *Foxg1* compared to pTa1-driven transgene (**Fig. 3DE**), this scenario suggests that Foxg1 down-regulation of *L1*-mRNA detectable in mid-neuronogenic cultures may have largely taken place in NPs (**Fig. 7** (2)).

Altogether these data point to a negative impact of Foxg1 on *L1*-mRNA expression, both in NPs and Ns. Moreover, mirror phenotypes displayed by *Foxg1*-LOF and -OE cultures further suggest that Foxg1 *physiologically tunes* these levels.

### Mechanisms underlying Foxg1 control of *L1* transcription

Foxg1 is known to often act as a transcriptional repressor [5,24]. We wondered whether this also specifically applies to *L1*s. To this aim we set early neuronogenic cultures, "wild type" (*Plap*-OE) and *Foxg1*-OE, and we evaluated Foxg1-enrichment at their *L1* loci against IgG controls, by chromatin immuno-precipitation (ChIP)-qPCR. We found that this enrichment was barely detectable in *Plap*-OE cultures and, conversely, statistically significant at all diagnostic amplicons in *Foxg1*-OE preparations (*p_5’UTR.A_*<0.014, *p_5’UTR.Gf_*<0.01, *p_5’UTR.Tf_*<0.002, *p_orf2_*<0.02, *p_3’UTR_*<0.01, with n=4,4) (**Fig. 6A**). Taking into account the prevalence of NSCs in early *Plap*-OE cultures and their massive conversion into NPs elicited by *Foxg1*-OE (**Fig. 5A** (1)), ChIP results shown in **Fig. 6A** suggest that Foxg1 recruitment at L1 loci can be negligible in NSCs and significant in NPs (**Fig. 7** (1)).

**Figure 6.**
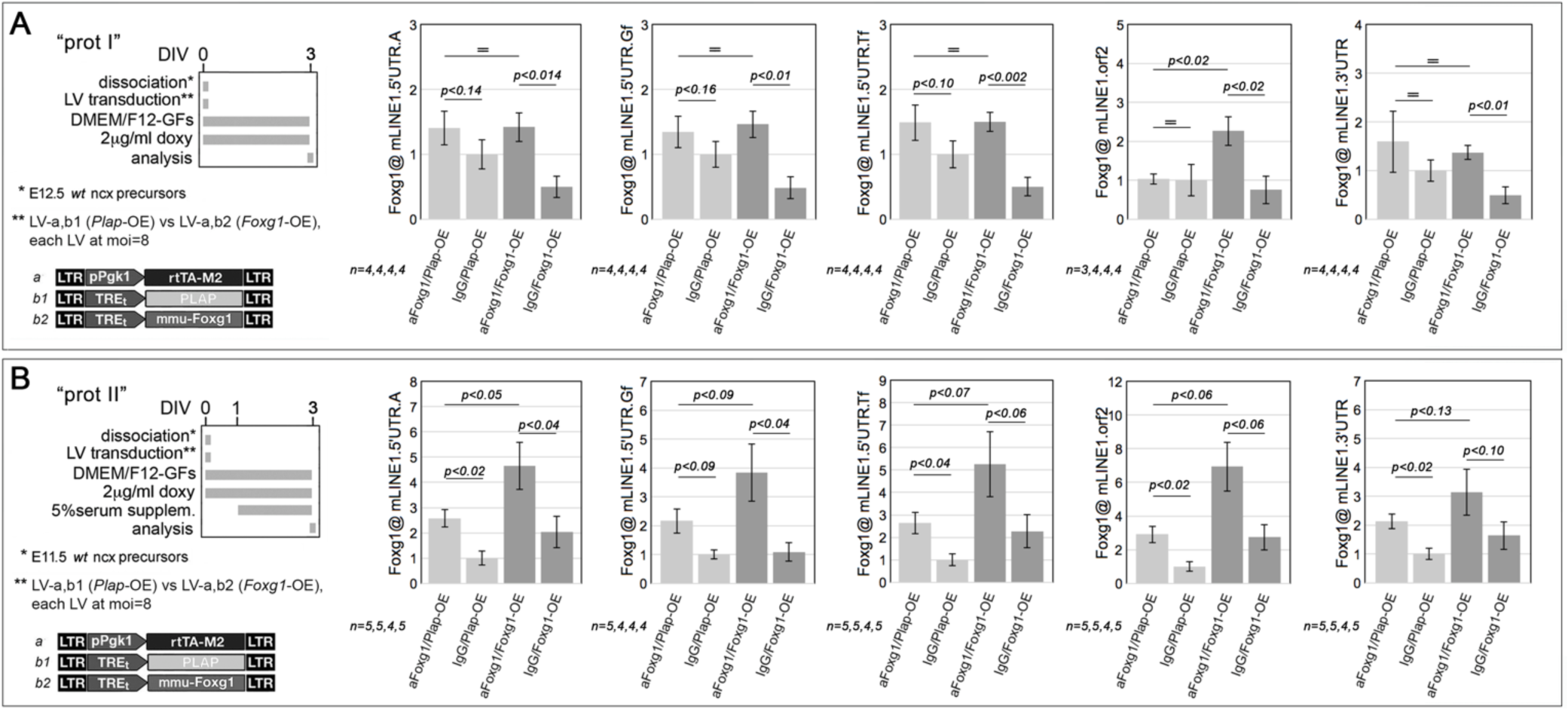
Chromatin immunoprecipitation (ChIP) profiling of Foxg1 protein enrichment at L1 elements. To left, protocols and lentiviral vectors employed, to right, results. Analysis run in early-(**A**) and mid-(**B**) neuronogenic cultures, set according to type I and II protocols, respectively. Cultures constitutively overexpressing *Foxg1* (or a *Plap* control), driven by the pPgk1 promoter. Chromatin immunoprecipitation performed by anti-Foxg1 antibody and control IgG. PCR quantitation of pan-*L1* diagnostic amplicons "L1.orf2" and "L1.3’UTR", and family-specific "L1.5’UTR" amplicons, ".A", ".Gf" and ".Tf". Results normalized against input chromatin. Scalebars, sem’s. Statistical evaluation of results by t-test (one-tail, unpaired). *n* = number of biological replicates, i.e. independently cultured and engineered cell aliquots originating from a common neural precursor pool.

**Figure 7.**
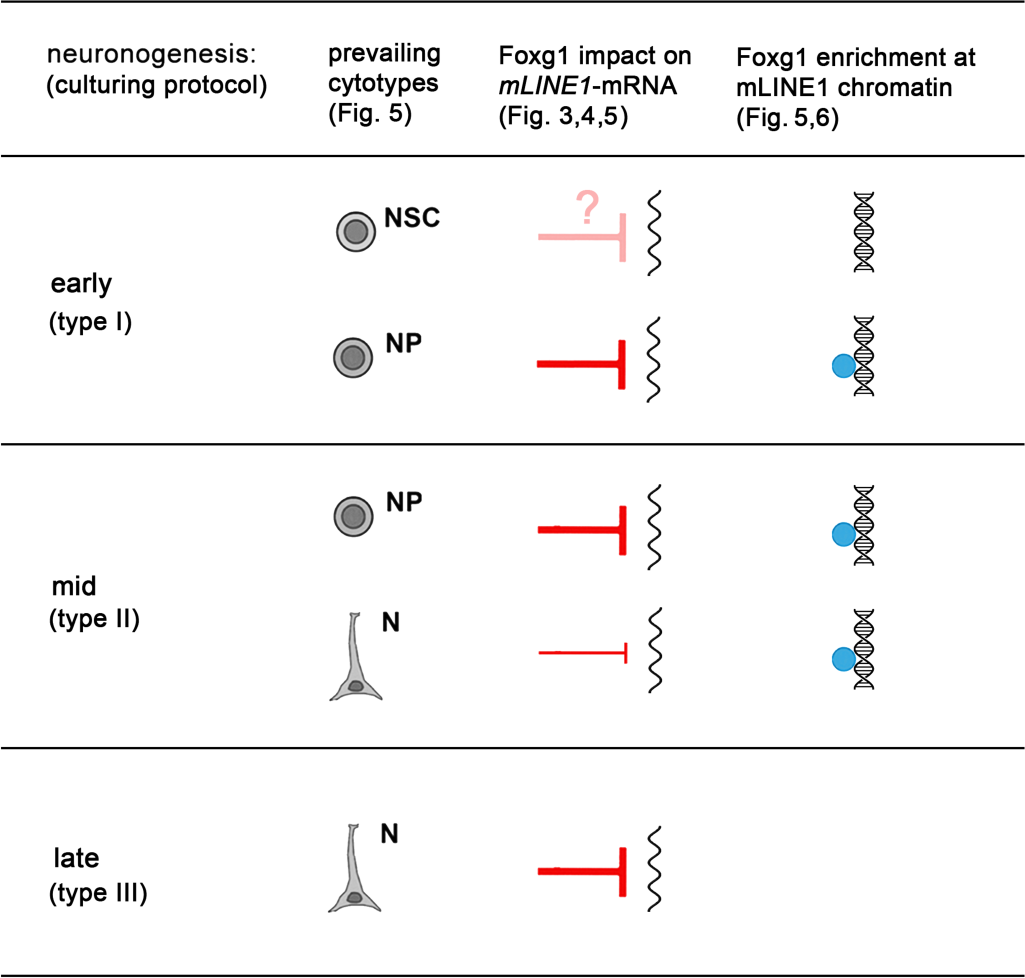
Synopsis of Foxg1 inhibition of *L1* expression within the pallial neuronogenic lineage. Tentative, temporal articulation of Foxg1 control of *L1* expression, based on integrated critical evaluation of L1-qRTPCR-(Fig. 3,4), Foxg1-qIF-(Fig. 5) and αFoxg1-ChIPqPCR-(Fig. 6) data.

Next, we run similar assays on chromatin of mid-neuronogenic cultures. In this case, a net Foxg1-enrichment was detectable at *all* diagnostic amplicons, regardless of "*Foxg1* genotype" of cultures, and such enrichment often reached formal statistical significance ("wild-type cultures": *p_5’UTR.A_*<0.02, *p_5’UTR.Gf_*<0.09, *p_5’UTR.Tf_*<0.04, *p_orf2_*<0.02, *p_3’UTR_*<0.02, with *n*=5,5 or - case Gf - *n*=5,4; *Foxg1*-OE cultures: *p_5’UTR.A_*<0.04, *p_5’UTR.Gf_*<0.04, *p_5’UTR.Tf_*<0.06, *p_orf2_*<0.06, *p_3’UTR_*<0.10, with n=4,5 or - case Gf - *n*=4,4) (**Fig. 6B**). Together with **Fig. 5C** (1) results, this scenario points to Foxg1 binding to *L1* loci in NPs and/or Ns (**Fig. 7**).

It has been shown that the dynamics of a number of epigenetic marks can be instrumental to tight control of *L1* transcription [52–59]. Moreover, inspection of the public Biogrid database [60] revealed that Foxg1 physically interacts with a number of effectors modulating the epigenetic landscape of chromatin, including Histone deacetylase 2 (HDAC2), Lysine-specific demethylase 5B (KDM5B), Lysine-specific demethylase 1A (KDM1A). Therefore, Foxg1 might exert its impact on *L1*-mRNA levels just by modulating the epigenetic state of *L1* chromatin. To address this issue, we scored chromatin extracted from from mid-neuronogenic cultures, *wild type* and *Foxg1*-OE, for its enrichment at *L1* loci for a number of key epigenetic markers: H3K4me3, H3K9me3, H3K27ac, MeCP2 (**Fig. 8A,B**). We found that H3K4me3, H3K9me3, H3K27ac were remarkably enriched at all diagnostic amplicons analyzed, *L1*.5’UTR.A, .Gf, .Tf, .orf2 and.3’UTR, regardless of culture genotype (about 60-1200-folds over controls). Conversely, MeCP2 enrichment over IgG controls was barely appreciable (<2-folds).

**Figure 8.**
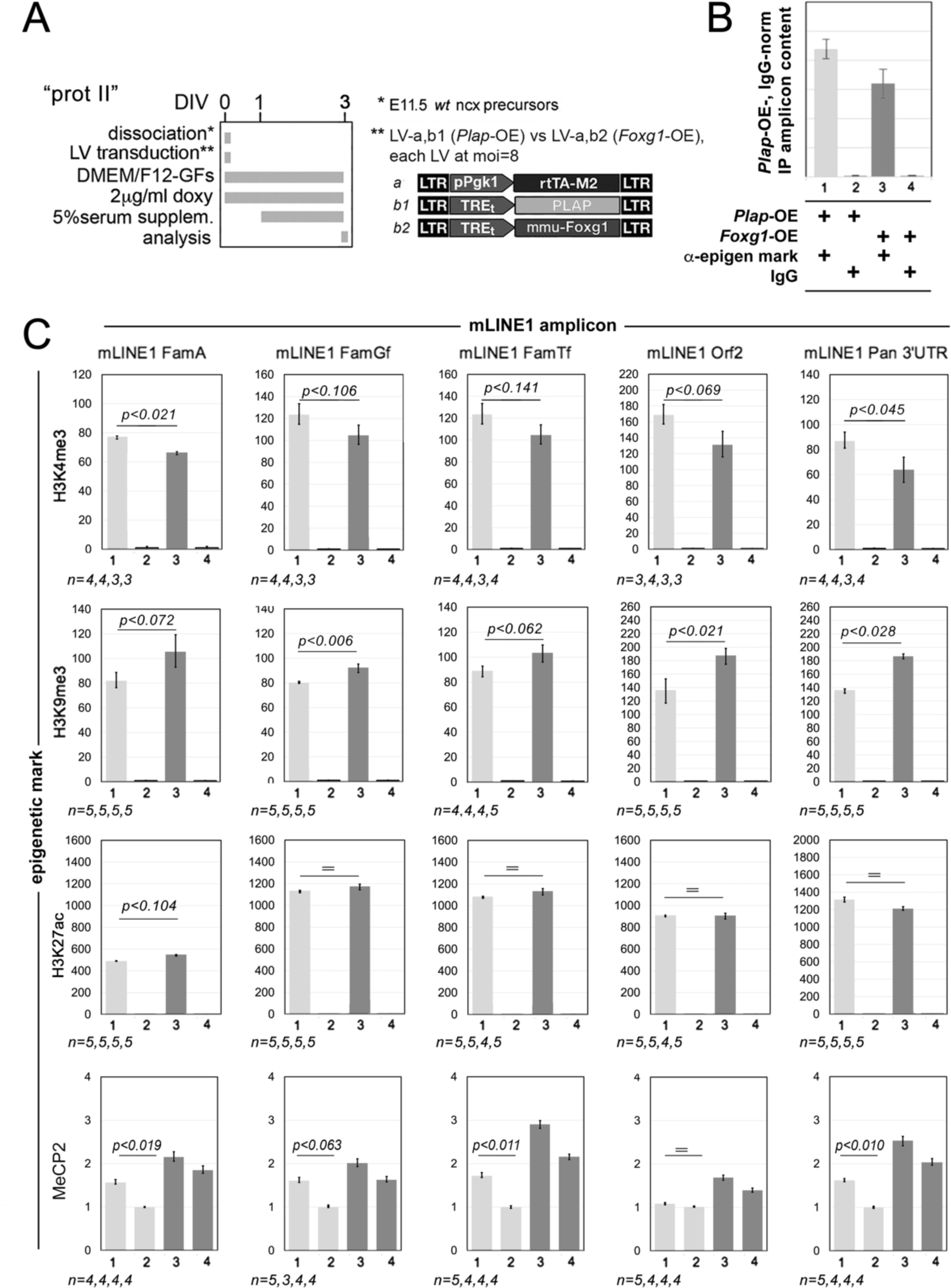
Chromatin immunoprecipitation (ChIP) profiling of H3K4me3, H3K9me3, H3K27ac, and MeCP2 enrichment at *L1* elements. In (**A**), protocols and lentiviral vectors employed, in (**B**) an illustrative prototype of results presentation, in (**C**) results. Analyses run in mid-neuronogenic cultures, set according to a type II protocol. Cultures constitutively overexpressing *Foxg1* (or a *Plap* control), driven by the pPgk1 promoter. Chromatin immunoprecipitation performed by αH3K4me3, αH3K9me3, αH3K27ac, and αMeCP2 antibodies and their isotypic IgG controls. PCR quantitation of pan-*L1* diagnostic amplicons "*L1*.orf2" and "*L1*.3’UTR", and family-specific "L1.5’UTR amplicons", ".A", ".Gf" and ".Tf". Results normalized against input chromatin. Scalebars, sem’s. Statistical evaluation of results by t-test (one-tail, unpaired). *n* = number of biological replicates, i.e. independently cultured and engineered cell aliquots originating from a common neural precursor pool.

Next, H3K4me3 enrichment was systematically reduced in *Foxg1*-OE compared to *Plap*-OE cultures (*p_5’UTR.A_*<0.021, *p_5’UTR.Gf_*<0.106, *p_5’UTR.Tf_*<0.141, *p_orf2_*<0.069, *p_3’UTR_*<0.045, with *n*=4,3 and - in case of orf2 - n=3,3). Conversely, an opposite dynamic could be detected in case of H3K9me3 (*p_5’UTR.A_*<0.072, *p_5’UTR.Gf_*<0.006, *p_5’UTR.Tf_*<0.062, *p_orf2_*<0.021, *p_3’UTR_*<0.028, with *n*=5,5 and - in case of Tf - n=4,4)(**Fig. 8C**).

Altogether, these results are consistent with the hypothesis that Foxg1 modulation of *L1* transcription is mediated by pervasive changes in the epigenetic state of these elements, namely a decrease of transcription-promoting H3K4me3 marks and an increase of heterochomatic H3K9me3 ones. On the other side, (1) high H3K27ac levels detectable in both controls and *Foxg1*-OE samples point to a transient bivalent state of chromatine, amenable to both silencing and transcription [55], and (2) low *L1* enrichment for MeCP2 might reflect relatively low expression of this protein at mid-neuronogenic stages [61].

Finally, to shed further light on mechanisms mediating Foxg1 impact on *L1* transcription, we took advantage of the neuropathogenic *FOXG1^W308X^* allele [6], encoding for a prematurely truncated protein, missing binding domains for the Groucho/Tle co-repressor and the *KDM5B*-encoded JARID1B H3K4me2/3-demethylase (**Fig. 3D**). Delivered to mid-neuronogenic cultures as a Tet^ON^-driven transgene, *FOXG1^W308X^* reduced *L1*-mRNA levels by 19.23±3.30% (*p*<0.006; *n*=5,5), 14.46±4.32% (*p*<0.013; *n*=5,5), and 8.40±2.59% (*p*<0.061; *n*=5,5), as evaluated at diagnostic amplicons "orf2", "5’UTR.A" and "5’UTR.Tf", respectively (**Fig. 9**). This down-regulation was far less pronounced compared to the that elicited by wild-type Foxg1 (**Fig. 3D**), suggesting JARID1B and/or Groucho/Tle to contribute to *Foxg1*-dependent *L1* repression.

**Figure 9.**
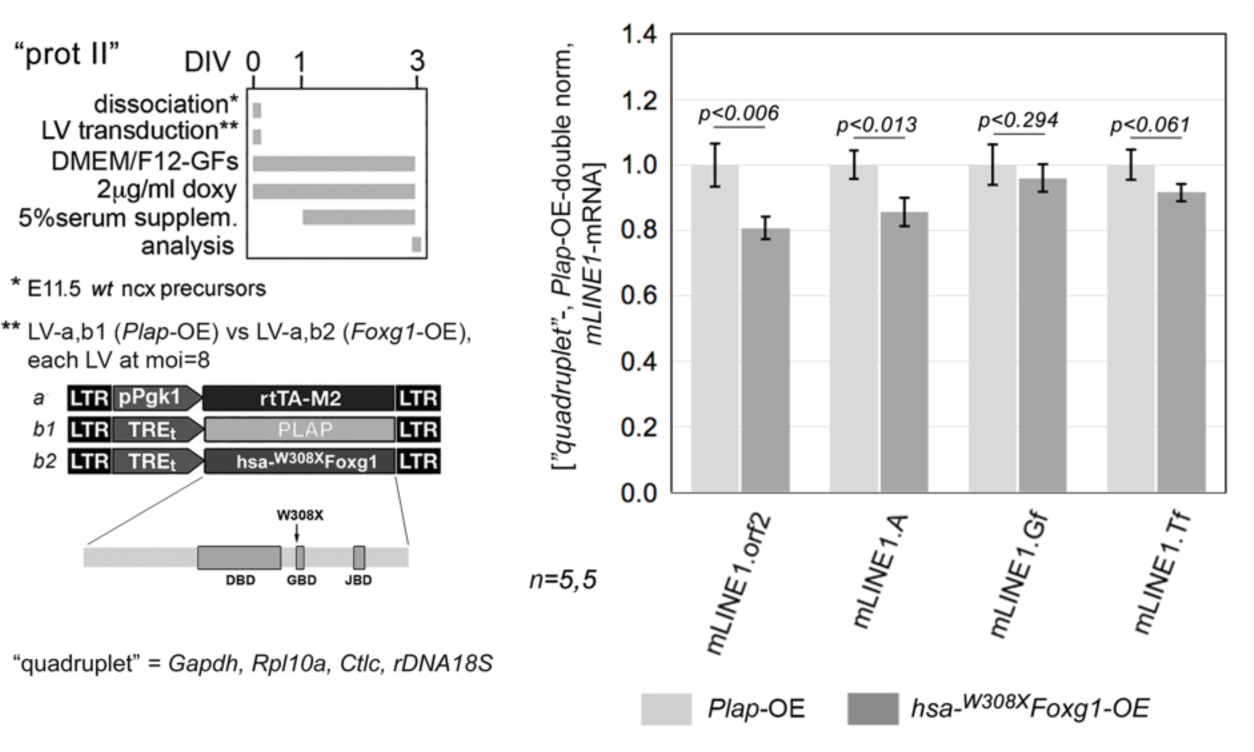
Modulation of *L1*-mRNA levels in murine mid-neuronogenic pallial cultures overexpressing the *hsa-^W308X^Foxg1* mutant allele. To left, protocol and lentiviral vectors employed. Transgenes driven by the constitutively firing pPgk1 promoter. The human (hsa) mutant allele in order encodes for a prematurely truncated protein, including the DNA-binding domain (DBD), but not the Groucho- and Jarid-binding domains (GBD and JBD, respectively). RT-PCR quantitation of pan-*L1* diagnostic amplicon "*L1*.orf2", and family-specific amplicons, "*L1*.A", "*L1*.Gf" and " *L1*.Tf", in neural cultures set according to a type II protocol. Data double normalized against gene quadruplet (*Gapdh*, *Rpl10a*, *Cltc* and *rDNA 18S*) and control values. Error-bars representing sem’s. Statistical significance of results evaluated by t-test (one-tailed, unpaired). *n* is the number of biological replicates, i.e. independently cultured and engineered cell aliquots, originating from pooled, *wild-type* E11.5 neocortical primordia.

### Temporal progression of pallial *L1* DNA copy-number

We wondered if, in addition to inhibiting *L1* transcription of *L1* elements, Foxg1 might further impact their DNA copy number. To get preliminary information about natural dynamics of *L1* DNA within the developing embryonic pallium, we scored early-, mid- and late-neuronogenic cultures for their cumulative *L1* copy number (**Fig. 10A**). We found that this number did not change across early- and mid-neuronogenic cultures, whereas it was increased by 35.08±3.65% (*p*<0.001, *n*=5.7) in late-neuronogenic ones (**Fig. 10B**, (1)). Actually, these results were obtained on DNA prepared by a dedicated sample digestion procedure ("high PK"), aimed at extracting DNA with comparable efficacy regardless of the compaction state of chromatin. [Fulfillment of this requirement had been previously tested, by quantifying an X-chromosomal/*lyonizable*, *Mecp2* amplicon in DNA extracted from female and male tissues, and normalizing it against an autosomal amplicon (*Gfap)*. This gave a normalized, female-to-male *Mecp2* signal ratio, equalling 1.54±0.18, with *p****_♂-♁_***<0.053, *n*=3,2 (**Fig. S2**(1))]. Next, to strenghten **Fig. 10B**, (1) results, we repeated the quantification of pallial *L1* content upon replacing the "high PK" protocol with a further improved version of it ("very high PK"). [With this latter protocol the female-to-male *Mecp2* signal ratio arose to 2.25±0.45 (*p****_♂-♁_***<0.016, *n*=3,2) and a similar 2.22±0.10 ratio was also obtained for the X-chromosomal *Cdkl5* DNA (*p****_♂-♁_***<0.001, *n*=3,3) (**Fig. S2**(2,3)]. Moreover, as a control, we included in this last assay late-neuronogenic cultures pre-treated by chronic lamivudine, an established inhibitor of retro-transcription. "Very high PK" samples substantially replicated the outcome of "high PK" ones, with *L1* copy number increased in late-neuronogenic cultures by 1.31±0.16-folds compared to their mid-neuronogenic counterparts (*p*<0.04, *n*=8,8). Remarkably, this increase was fully suppressed by lamivudine (**Fig. 10B**(2)). As somatic retro-transcription has been reported to generate new *L1* copies missing the 5’UTR, assays referred to in **Fig. 10B**(1,2) relied on the "3’UTR" diagnostic amplicon. In this respect, as a further control, we also compared mid- and late neuronogenic cultures for their L1 content by means of sub-family-specific 5’UTR amplicons. As expected, no relevant changes were found (**Fig. 10B**(3,4,5), except an increasing trend at the "5’UTR.A" amplicon **(Fig. 10B**(3)), possibly reflecting differential somatic RT failure in distinct mLINE sub-families.

**Figure 10.**
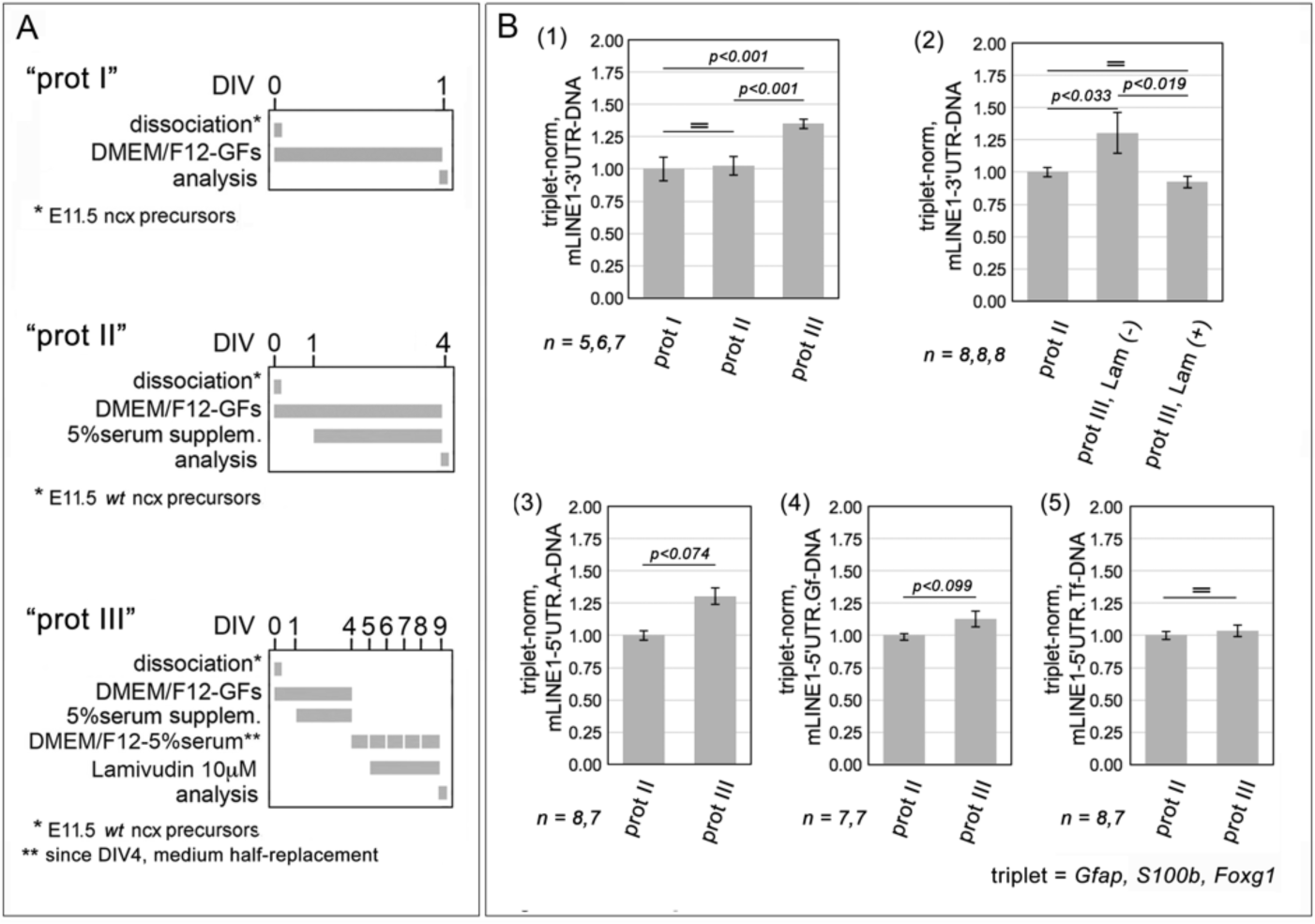
*L1* DNA copy number variations (CNVs) in early-mid- and late-neuronogenic murine pallial cultures. **(A)** protocols, **(B)** results. Early-mid- and late-neuronogenic cultures set by means of type I, II and III protocols, respectively. DNA extraction performed by "high PK" (1) and "very high PK" procedure (2-4) (see Materials and Methods). DNA CNVs assessed by quantitative PCR, followed by normalization against endogenous *Gfap*, *S100b* and *Foxg1* (gene "triplet"). In graph (1), shown is the total *L1* copy number detectable in late- and mid-normalized against early-neuronogenic cultures. In graph (2), shown is the suppression of *L1* copy number variation which normally occurs between "mid-" and "late-neuronogenic" cultures, elicited by the pan-RT inhibitor lamivudine (aka 3T3). Finally, in graphs (3-5), as controls, shown are comparisons of family-specific, DNA copy numbers, as detected by "5’UTR.A", "5’UTR.Gf" and "5’UTR.Tf" diagnostic amplicons. Error-bars representing sem’s. Statistical significance of results evaluated by t-test (one-tailed, unpaired). *n* is the number of biological replicates, i.e. independently cultured and engineered cell aliquots, originating from pooled, *wild-type* E11.5 neocortical primordia.

Finally, to validate the dynamics of *mLINE*-DNA observed in mid-vs late-neuronogenic cultures, we compared *mLINE*-DNA content in neocortices dissected from E14.5 vs P0 wild-type mice. As expected, the latter exceeded the former, by 23.8±4.7% (p<0.006; n=8,7) (**Fig. 11**), corroborating our previous findings. Intriguingly, an increase of *mLINE*-DNA content over the same time interval was also detectable in the mesencephalic tectum, where its amplitude was even larger (+54.0±8.1%, with p<0.001 and n=6,9) (**Fig. 11**).

**Figure 11.**
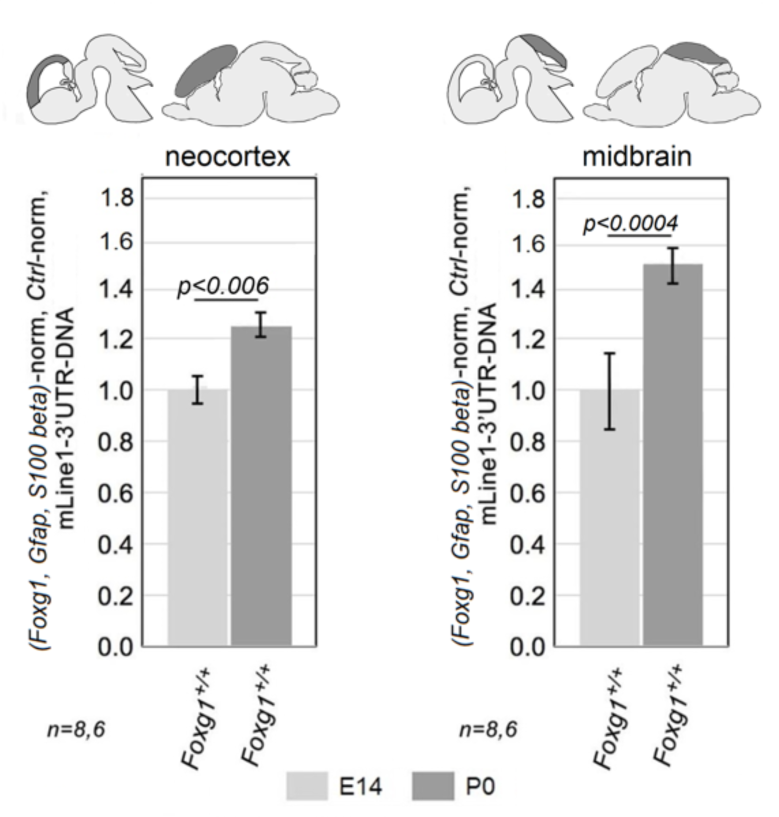
*L1* DNA copy numbers in neocortex and tectum of wild type, mid-neuronogenic and perinatal mice. PCR quantitation of the pan-*L1* diagnostic amplicon "*L1*.3’UTR". Data normalized against the *Foxg1/Gfap/Nfia* gene triplet. Error-bars representing sem’s. Statistical significance of results evaluated by t-test (one-tailed, unpaired). *n* is the number of biological replicates, i.e. neocortices taken from distinct pups.

### Foxg1 impact on *L1*.DNA copy numbers

We have shown that *L1* copy number increases during neocortical neuronogenesis progression. To further investigate the role (if any) of *Foxg1* in this process, we compared *L1* DNA content in neocortices of *Foxg1^−/+^* neonates and their littermate wild type controls. Normalized against *Gfap* and *Nfia,* such content turned out to be decreased in *Foxg1*-LOF samples by 7.50±0.85% (*p*<10^−3^, *n*=6,8), compared to controls (**Fig. 12**). To note, this variation approximately equals 1/3 of the increment in neocortical *L1* copies detectable over the same time interval in wild type mice (**Fig. 11**). Moreover, it occurred in mutants characterized by *Foxg1*-mRNA levels reduced by only 33.59±6.17% (*p*<0.02, *n*=8,7) compared to wild-type controls (**Table S1**).

**Figure 12.**
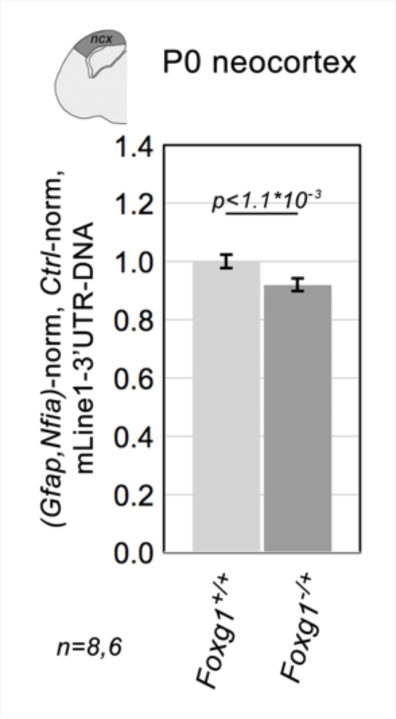
*L1* DNA copy numbers in neocortex of *Foxg1^−/+^* mouse neonates and controls. PCR quantitation of the pan-*L1* diagnostic amplicon "*L1*.3’UTR". Data double normalized against the *Gfap/Nfia* gene doublet, and wild-type controls. Error-bars representing sem’s. Statistical significance of results evaluated by t-test (two-tailed, unpaired). *n* is the number of biological replicates, i.e. neocortices taken from distinct pups.

To corroborate these findings, we repeated *L1* copy number evaluation in primary, late-neuronogenic cultures manipulated by CRISPR-Cas9 technology, which allowed us to achieve a more pronounced, -65.37±2.68% (*p*<10^−7^, *n*=8,8) *Foxg1* down-regulation, (**Table S1**). Remarkably, in this case, *L1* copy number was reduced by 15.23±2.19% (upon normalization against *Gfap* and *Nfia*, with *p*<10^−3^, *n*=8,8) (**Fig. 13B**(1)), corresponding to about 2/3 of the "physiological" increment referred to above (**Fig. 11**).

**Figure 13.**
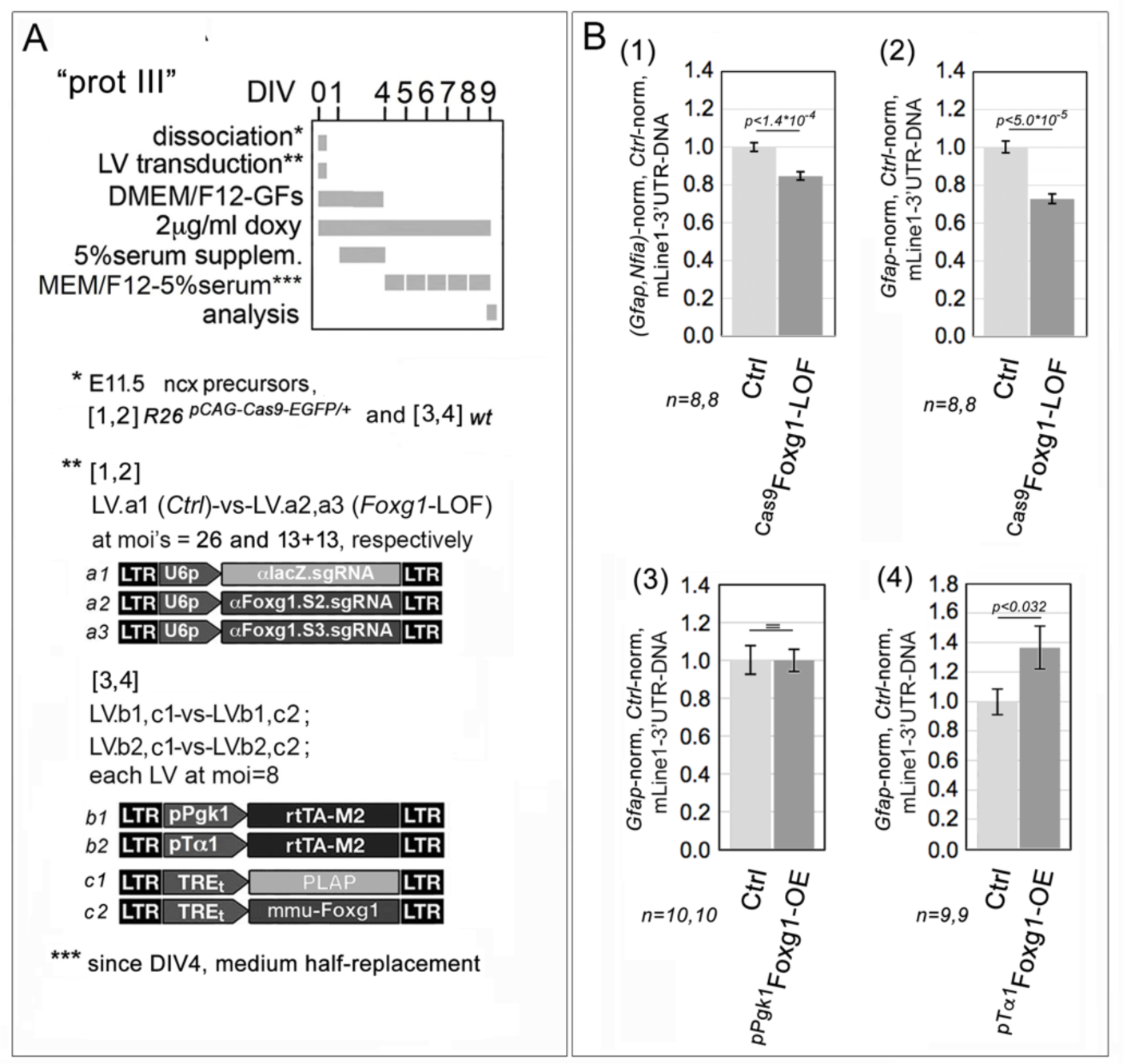
Impact of *Foxg1*-knock-down (KD) and *Foxg1*-OE on *L1* DNA copy numbers in late-neuronogenic pallial cultures. In (**A**), protocols (with lentiviruses employed), in (**B**) results. CRISPR-Cas9 machinery driven by constitutively active U6p (sgRNAs) and R26/pCAG (Cas9) promoters, Foxg1 transgene led by constitutive Pgk1p and NP/N-restricted pTa1. Neural cultures set according to a "type III" protocol. Assesment of total L1 copies by quantitation of the "*mLINE*.3’UTR" amplicon. Data double-normalized, against *Gfap* (or the *Gfap/Nfia* gene doublet) and control values. Error-bars representing sem’s. Statistical significance of results evaluated by t-test (one-tailed, unpaired). *n* is the number of biological replicates, i.e. independently cultured and engineered cell aliquots, originating from pooled, *R26^pCAG-Cas9-EGFP/+^* (1,2) and *wild-type* (3,4), E11.5 neocortical primordia.

To achieve a more comprehensive picture of Foxg1 role in tuning *L1* copy number, we finally overexpressed it in late-neuronogenic cultures, under the control of pTa1 and pPgk1 promoters, and we evaluated the impact of these manipulations on *L1* DNA content. As expected, we found that such content was increased by "pTa1-driven *Foxg1",* by 37.11±16.35%, with *p*<0.03, and *n*=9,9 (**Fig. 13B**(4)). Conversely, no *L1*-DNA increase was elicited by "pPgk1-driven *Foxg1"* (**Fig. 13B**(3)), possibly due to the stronger *L1*-mRNA downregulation triggered by such transgene compared to its pTa1 counterpart (**Fig. 3D,E**). Altogether, these results point to Foxg1 as a key player needed for *L1* DNA amplification and *its fine tuning*. To note, such amplification-promoting activity has to be particularly robust, as it emerged despite the concomitant, *Foxg1*-induced down-regulation of *L1*-mRNA, namely the *starting template* from which new *L1* DNA is generated.

### Mechanisms underlying Foxg1 impact on *L1* copy number

We wondered: how does Foxg1 impact *L1* DNA content? We considered two possible scenarios: (1) it acts indirectly, as a "professional transcription factor" modulating transcription of key effectors involved in synthesis and/or degradation of new somatic *L1* copies; (2) it straightly regulates these processes, through physical interaction with factors implicated in them and/or *L1*-mRNA.

As for (1), we inspected a database of genes mis-regulated upon Foxg1 overepression in neocortical neuronal cultures [51]. We found that mRNA encoding for Apobec1, an inhibitor of *L1* retro-transposition [62], is halved in *Foxg1*-OE samples (**Table S3A**), suggesting Foxg1 might promote retro-transposition, mitigating such inhibition.

As for (2), we interrogated the public Biogrid database for Foxg1 interactors implicated in retro-transposition control and we found two well known antagonizers of *L1* retro-transposition, Mov10 and Ddx39a [63](**Table S3B**). We co-manipulated expression levels of each of them and *Foxg1* in late-neuronogenic cell preparations, and we evaluated the impact of such interventions on *L1*-DNA content (**Fig. 14A**). Interestingly, in a sensitized *Foxg1*-lof environment, "wild-type" levels of both *Mov10* and *Ddx39a* gave rise to statistically significant decreases in *L1*-DNA copy number compared to their knock-down counterparts [−16±3% with p<0.035 n=7, as well as -18±3% with p<0.018, and n=6, respectively]. Conversely, in a *Foxg1*-wt environment, a decrease was only detectable in *Ddx39a* "wild type" compared to *Ddx39a* knock-down samples [−15±3%, with p<0.015, and n=10]. Intringuingly, two-ways ANOVA analysis of results gave signals of *statistical* interaction among *Foxg1* on one side and *Mov10* (p<0.026) or *Ddx39a* (p<0.058) variables on the other (**Fig. 14B**). All that points to a likely *functional* interaction taking place among Foxg1 and the helicases encoded by these two genes.

**Figure 14.**
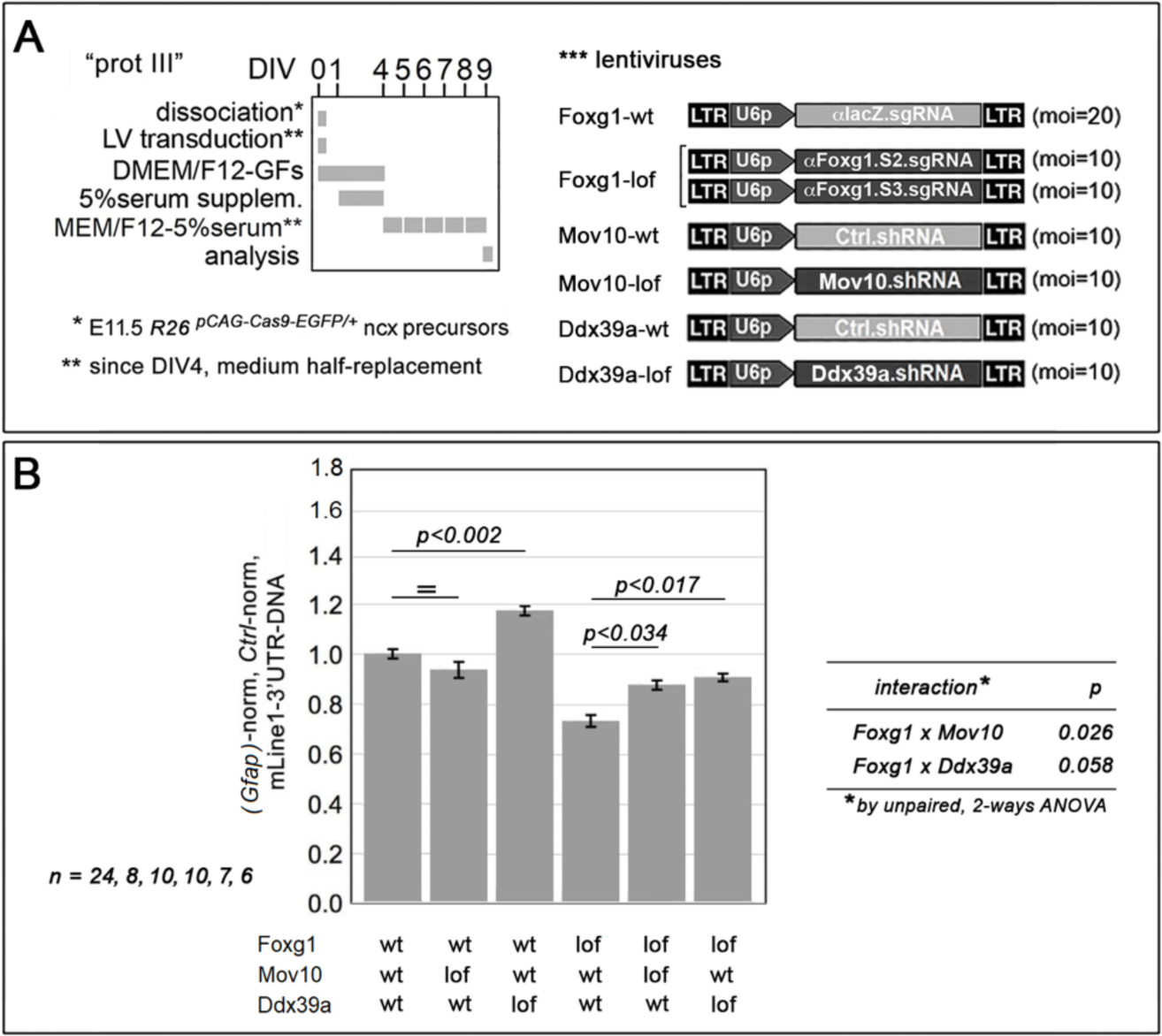
Functional interaction among *Foxg1* and helicase genes *Mov10* and Ddx39a in modulation of L1 copy number. In (**A**), protocols (with lentiviruses employed). In (**B**) shown is the impact exerted by *Mov10* and *Ddx39a* downregulation, in *Foxg1*-wt and *Foxg1*-LOF environments, on L1 copy number, as evaluated in late-neuronogenic cultures by qPCR. Data double-normalized, against *Gfap* and control values. n is the number of biological replicates, i.e. independently cultured and engineered preparations, originating from a common neural cell pool. Statistical evaluation of results performed by t-test, one-tailed and unpaired, as well as by, unpaired, 2-ways ANOVA. Errors bars indicate s.e.m.

Next, we reasoned that Foxg1 could counteract Mov10 and Ddx39a, by preventing them from interacting with *L1*-mRNA. This might be achieved by chelating the helicase in order and/or shielding its *L1*-RNA interactor. As said, the former phenomenon has been already documented [64]. To assess the latter, we run a set of RNA-immunoprecipitation (RIP) assays, by which we quantified "5’UTR", "orf2" and "3’UTR" *L1* diagnostic amplicons in anti-Foxg1-immunoprecipitated RNA (**Fig. 15**). Consistently with our prediction, we found a robust enrichment of Foxg1 at both 5’ and 3’ ends of L1-mRNA. Referring to IgG controls, this equalled 3.11±0.20 folds at "5’UTR.A" (p<10^−4^, n=5,3), 1.85±0.28 folds at "5’UTR.Gf" (p<0.043, n=5,3), 4.13±0.51 folds at "5’UTR.Tf" (p<0.003, n=5,3), and 3.89±0.55 folds at "3’UTR" (**Fig. 15** (1,2,3,5)). Foxg1 enrichment was lower at "orf2", where it did not reach statistical significance (**Fig. 15** (4)).

**Figure 15.**
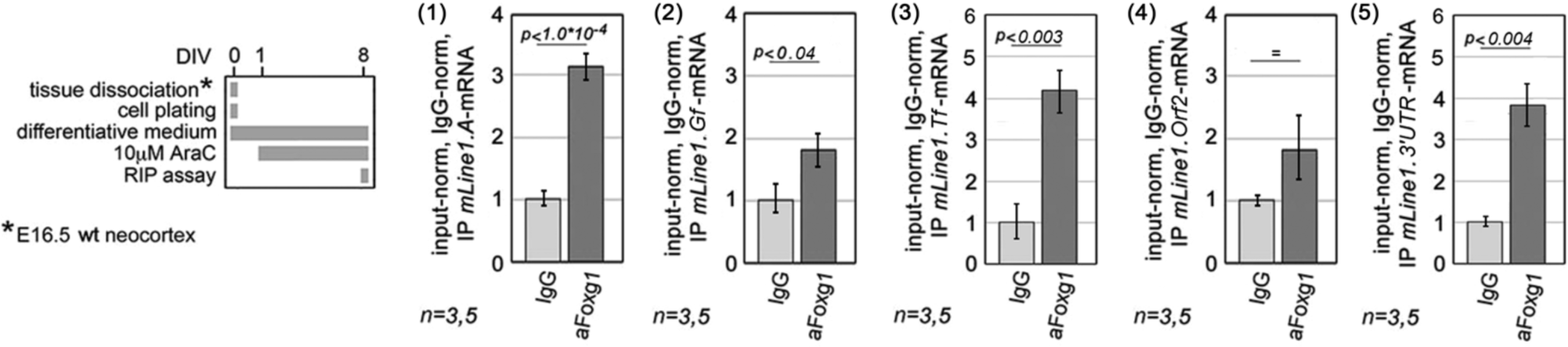
Foxg1-protein/*L1*-mRNA interaction in neocortical neurons, as evaluated by RNA immunoprecipitation-quantitative PCR (qRIP-PCR) assay. To left, protocol, to right, results. Diagnostic amplicons, L1.5’UTR.A (1), L1.5’UTR.Gf (2), L1.5’UTR.Tf (3), L1.orf2 (4), and L1.3’UTR (5). Results double normalized, against input-RNA and IgG-IP samples. Througout figure, n is the number of biological replicates, i.e. independently cultured and engineered preparations, originating from a common neural cell pool. Statistical evaluation of results performed by t-test, one-tailed and unpaired. Errors bars indicate s.e.m.

## DISCUSSION

Foxg1 masters telencephalic development, exerting a highly pleiotropic control over it. *L1* is a prominent retrotransposon family likely contributing to plasticity of neuronal genomic DNA. Here we systematically investigated the impact of the former on the biology of the latter in the developing murine neocortex. Main results were as follows.

As suspected, we found that *L1*-mRNA encoded by all three retro-transposition-competent sub-families (A, Gf and Tf) was increased in *Foxg1*-LOF mouse neonates compared to wild type controls (**Fig. 1**). To model temporal and lineage articulation of *Foxg1*-dependent *L1* regulation, we developed an integrated culture-set, representing early-, mid- and late-phases of neuronogenesis in vitro. This set showed an increasing trend of *L1* expression paralleling in vivo *L1*-mRNA progression (**Fig. 2**). Next, we manipulated Foxg1 levels at different stages of the neuronogenic progression, by means of distinctive neural cell type-specific promoters (**Fig. 3,4**). We evaluated the impact of these manipulations on sizes of NSCs, NPs and Ns compartments, and we mapped changes of Foxg1 protein levels to these compartments (**Fig. 5**). Moreover, we profiled Foxg1 binding to *L1* chromatine at different steps of neuronogenic progression (**Fig. 6**). Integrated analysis of **Fig. 2-6** results provided us robust evidence that Foxg1 represses *L1* expression, selectively in NPs and Ns (**Fig. 7**). As expected, we found that such repression was associated to reduced H3K4me3 and increased H3K9me3 marks along the entire retrotransposon (**Fig. 8**). A prematurely truncated, human neuropathogenic, loss-of-function variant of FOXG1 downregulated *L1*-mRNA too, although weaker than its "healthy" counterpart (**Fig. 9**).

Propedeutically to investigating a possible impact of *Foxg1* on *mLINE*-DNA copy number, we profiled progression of this number in early-, mid- and late-neuronogenic cultures, and we found that it was increased by about 35% in late ones, in a retro-transcription-dependent way (**Fig. 10**). A comparable increase was found in vivo as well, in neonatal compared to mid-neuronogenic embryonic neocortex, suggesting the former phenomenon to be genuine (**Fig. 11**). Next, unexpectedly, we discovered that Foxg1 down-regulation, in vivo as well as in vitro, elicited a remarkable, Foxg1-dose-dependent reduction in *L1*-DNA content, up to 2/3 of the natural increase the parameter would undergo in vivo. Consistently, mild Foxg1 overexpression in mid-neuronogenic cultures increased *L1*-DNA, further suggesting that Foxg1 *tunes* physiological amplification of such DNA (**Fig. 12,13**). Hypothesizing a straight intervention of Foxg1 in *L1* retrotranscription, we evaluated the outcome of combinatorial manipulation of Foxg1 and two interactors of it known to inhibit retrotransposition, Mov10 and Ddx39a. As expected, we found that Foxg1 desensitizes neocortical neurons to the activity of these helicases, resulting in increased L1-DNA copy number (**Fig. 14**). Finally, consistently with the hypothesis mentioned above, we found that Foxg1 binds to *L1*-mRNA, with special emphasis on its 5’ and 3’ ends (**Fig. 15**).

In synthesis, two main messages emerged from our study

First, we demonstrated that Foxg1 robustly inhibits pallial *L1*-mRNA expression (**Fig. 3,4**), by binding *L1* chromatin (**Fig. 6**) and exerting a deep impact on its epigenetic state (**Fig. 8**). Trans-repression was detectable in case of all three retrotransposition-competent L1 sub-families, albeit the Gf one turned out to be the most sensitive to Foxg1 manupulation, both up- and down-wards (**Fig. 1,3**). In this way, Foxg1 adds to the small TF-set shown to control *L1* transcription in CNS [65–70]. In this context, it recalls Sox2. However, Sox2 was documented to represses L1 transcription in adult NSCs [66], while Foxg1 has been shown to act in embryonic neuronal progenitors and neurons (**Fig. 7**). Moreover, Sox2 is expressed in the apical compartment of the whole neuraxis [71], Foxg1 is mainly confined to the telencephalon [72]. Actually, to our knowledge, Foxg1 is the first patterning gene proven to limit *L1* expression within a specific domain of the developing mouse brain [65,66].

Second, we documented an appreciable increase of pallial *L1*-DNA content occurring during late-intrauterine development, consistently with a previous report [46]. We discovered that, albeit reducing *L1*-mRNA (**Fig. 3**), an increase of Foxg1 levels around the baseline often leads to an augmented *L1*-DNA content (**Fig. 12,13** (1,2,4)). However, if the decline of *L1*-mRNA elicited by *Foxg1* overexpression is too pronounced (compare **Fig. 3D** vs **3E**), than the same genetic manipulation fails to increase *mLINE*-DNA content (compare **Fig. 13B**(3) vs **13B** (4)). In this respect, it is tempting to speculate that the transient down-regulation *Foxg1* physiologically undergoes in newborn pyramids [8] may be instrumental in allowing sufficient accumulation of *L1*-mRNA needed for subsequent retro-transcription. To note, the relationship between *L1*-mRNA and and *L1*-DNA levels depends on the CNS structure in order. Differently from neocortex, a huge amplification of *L1*-DNA content is achieved within the late-gestational tectum, despite the concomitant down-regulation of the "underlying" *L1*-mRNA level (**Fig. 2,11**).

Concerning *Foxg1* control of *L1*-DNA content, we hypothesized that it could take place via two helicases, Mov10 and Ddx39a, reported to antagonize *L1* retro-transcription [63] and physically interact with Foxg1 protein [64]. We rule out transcription as a mediator of this mechanism. In fact, while resulting in *Mov10*- and *Ddx39a*-mRNA downregulation, by -45.3% and -17.6%, respectively, with padj<0.05 (Artimagnella and Mallamaci, unpublished data), *Foxg1* knock-down did not increase *L1*-DNA content, but *reduced* it. Conversely, we noticed that *Foxg1* knock-down made the decline of *L1*-DNA evoked by higher levels of *Mov10* and *Ddx39a* more pronounced (**Fig. 14**). In addition, we showed Foxg1 protein also normally binds to *L1*-mRNA (**Fig. 15**). Hence, we propose that Foxg1 may ease *L1-*mRNA retro-transcription largely by preventing the interaction among the two helicases and such mRNA [63], competitively or because of steric hindrance. Such involvement in retrotranscription-control adds to Foxg1 implication in a number of other non-transcriptional metabolic routines, such as post-transcriptional ncRNA processing [25], translation [26] and mitochondrial biology [27].

We presently ignore the overall meaning of *Foxg1* control upon *L1* biology. On one side, *Foxg1* exerts a non-monotonic impact on neuronogenesis progression, (1) stimulating the NSC-to-NP transition ([5] **Fig. 5A**(1)), (2) inhibiting NPs exit from cell cycle [4], (3) promoting postmitotic neuronal differentiation [6,8,10–13], and (4) finally declining [13,73]. On the other side, it has been shown that specific ensembles of transposable elements are transcribed concomitantly with the progression of well-defined, early histogenetic routines [30] and, in some cases, it has been proven that such transcription is needed for the progression of these routines [31–33]. It is tempting to speculate this might apply to the neuronogenic lineage as well. In this context, *L1* trans-repression by Foxg1, in NPs and neurons, might be instrumental in mediating specific aspects of the control that this factor exerts on neuronogenesis progression. In addition, it has been proposed that somatic retrotransposition may help diversifying neuronal functional properties [36,74]. Upregulating *L1* copy number, Foxg1 might just promote this phenomenon.

Beyond its physiological articulation in the developing rodent embryo, the relationship between *FOXG1* and *L1* elements might be relevant to the etiopathogenesis of the human *FOXG1* syndrome. Deficient FOXG1 activity linked to *FOXG1* hemizygosity or heterozygosity for LOF-alleles might lead to *L1*-mRNA upregulation, supernumerary or GOF *FOXG1* alleles might result in exaggerated *L1*-DNA neo-synthesis. Both scenarios are of potential neuropathogenic interest [75] and, in this respect, early treatment with FDA-approved inhibitors of retrotranscription [76] might mitigate consequences of *FOXG1*-GOF mutations. However, major differences characterize cortical histogenesis and *L1* biology in humans and rodents. For these reasons, these issues definitively deserve further in depth investigations.

## MATERIALS AND METHODS

### Animal handling

Animal handling and subsequent procedures were in accordance with European and Italian laws [European Parliament and Council Directive of 22 September 2010 (2010/63/EU); Italian Government Decree of 4 March 2014, n° 26]. Experimental protocols were approved by SISSA OpBA (Institutional SISSA Committee for Animal Care).

Embryos and animals were generated at the SISSA mouse facility, as follows:

- wild-type ones were generated by breeding CD1 parents, purchased from Envigo Laboratories, Italy;
- *Foxg1*^+/-^ ones (and their wild type controls) were generated by breeding CD1-backcrossed, *Foxg1*^+/-^ males [49] to wild type CD1 females
- *Rosa26^pCAG-Cas9-2P2-Egfp)/+^* ones were generated by breeding *Rosa26^pCAG-Cas9-2P2-Egfp)/+^* males [originating from a line obtained by intercrossing a R*osa26^(pCAG-flSTOP-Cas9-2P2-Egfp)/+^* founder [77] to constitutive cre-expressors [78], and kept on a C57Bl/6 background] to wild type CD1 females.

Animals were staged by timed breeding and vaginal plug inspection. Where due, pregnant females were sacrificed by cervical dislocation.

As for genotyping:

- *Rosa26^pCAG-Cas9-2P2-Egfp)/+^* embryos were distinguished from their wild type littermates by inspection under fluorescence microscope
- *Foxg1^+/-^* embryos were distinguished from their wild type littermates by PCR genotyping, as previously described [3].

Molecular sexing was performed by a dedicated procedure, run in parallel with the microdissection of neural tissue of interest. For this purpose, a skin fragment from each embryo was collected and DNA extracted from it was used for fast, PCR-based genotyping. Males were distinguished by an oligo pair specifically amplifying the Y-chromosome-located *Uty* gene (see **Table 1**).

**Table 1.**
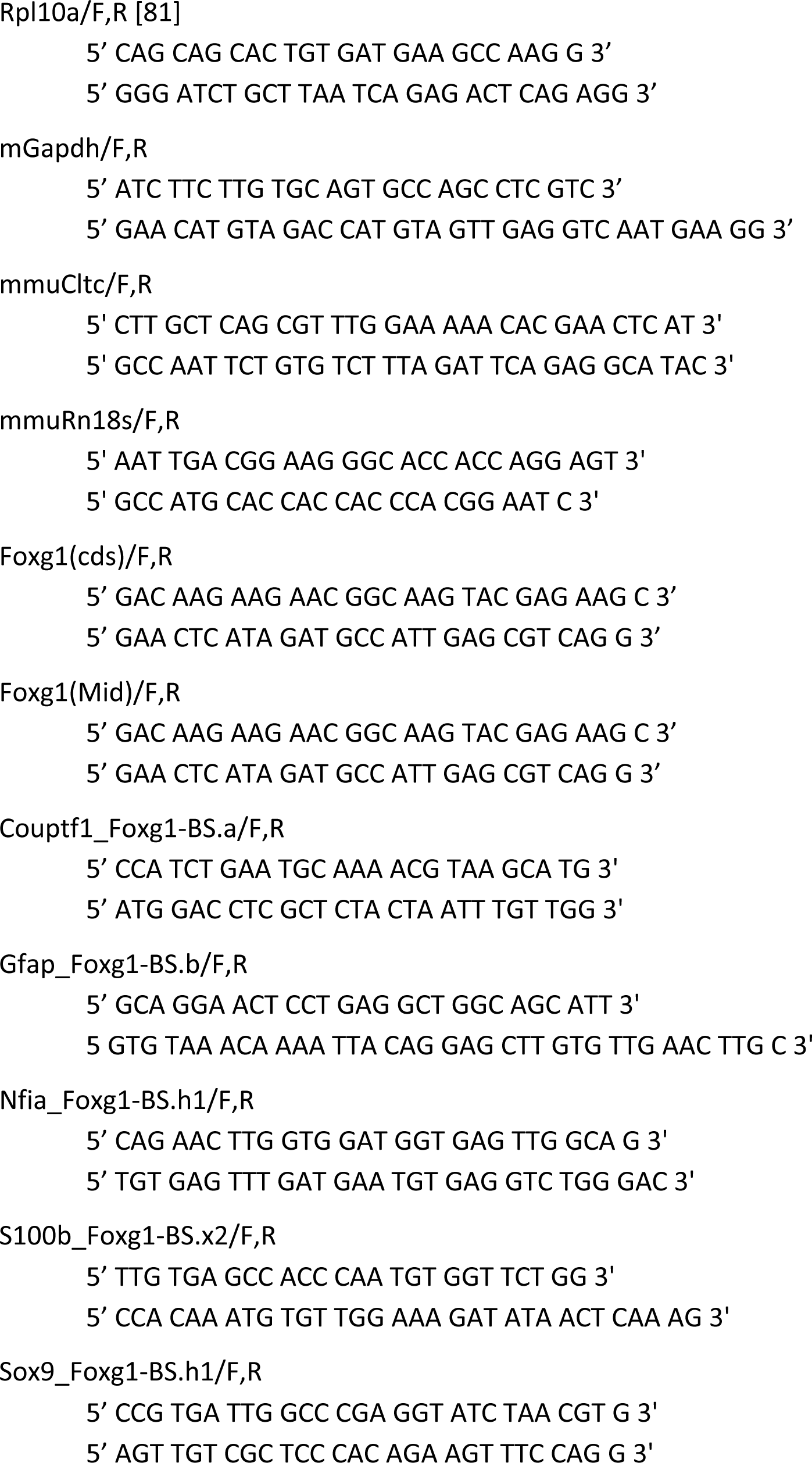
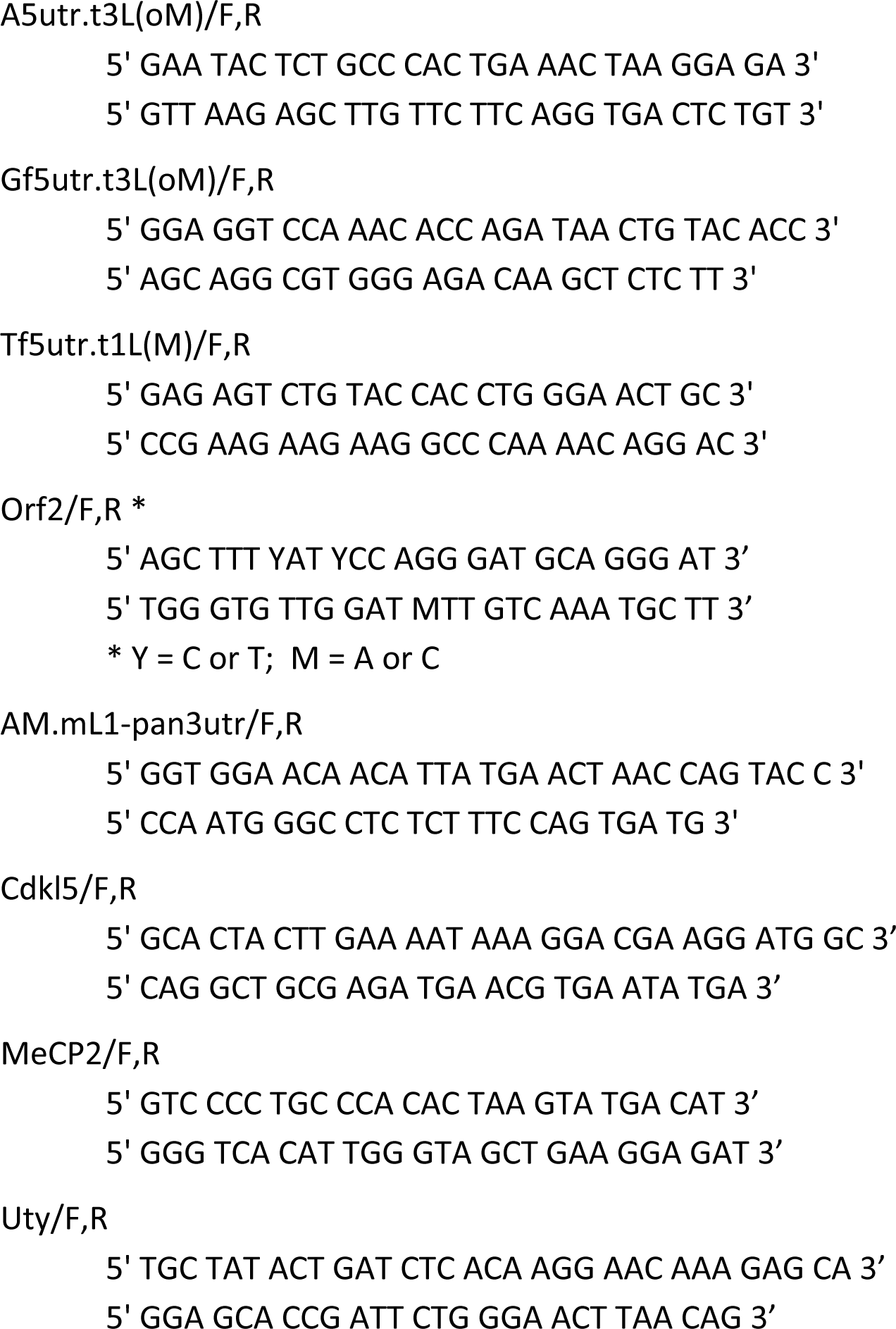
Primer sequences. The following oligonucleotide pairs have been employed in this study:

In general, extraction of genomic DNA employed for genotyping and preparation of the PCR reaction mix were performed by a KAPA HotStart Mouse Genotyping Kit Roche (KK7351), according to Manufacturer’s instructions.

### Primary neocortical cultures: early-, mid-, and late-neuronogenic ones

E11.5 - E12,5 mouse neocortical primordia were dissected and mechanically dissociated to single cells by gentle pipetting. Dissociated cells were quantified in a Burker chamber and then plated in 24-multiwell plates (Falcon) at 100 - 300 cells/uL, in pro-proliferative medium (1:1 DMEM-F12, 1X Glutamax (Gibco), 1X N2 supplement (Invitrogen), 1mg/ml BSA, 0.6% Glucose, 2 μg/ml mouse heparin (Stemcell technologies), 1X Pen/strept (Gibco), 10µg/ml Fungizone (Gibco), 20 ng/ml bFGF (invitrogen), 20 ng/mL EGF (Invitrogen). If required, neural cells were transduced with a LV mix, each LV at a multeplicity of infection (m.o.i.) ≥8, sufficient to infect almost the totality of neural cells in these conditions [4]. Neural cells were subsequently cultured according to three different schedules, aiming to model early, mid and late phases of neuronogenic progression:

<u>(1) Protocol I (Prot-I), early-neurogenic cultures</u> Already plated in pro-proliferative medium (see above) at 300 cells/µL, cells were kept in such medium up to DIV0.87-DIV3, and then processed for analysis.
<u>(2)  Protocol II (Prot-II), mid-neuronogenic cultures</u> Upon plating cells in pro-proliferative medium (see above) at 300 cells/µL, their medium was further supplemented with 5% FBS at DIV0.87. Cells were kept in the resulting medium up to DIV3, and then processed for analysis.
<u>(3)  Protocol III (Prot-III), late-neuronogenic cultures</u> Upon plating cells in pro-proliferative medium (see above) at a 100 cells/µL, their medium was further supplemented with 5% FBS at DIV0.87, and - next - daily hemi-replaced by "1:1 DMEM-F12, 1X Glutamax (Gibco), 1X N2 supplement (Invitrogen), 1mg/ml BSA, 0.6% Glucose, 2 μg/ml mouse heparin (Stemcell technologies), 1X Pen/strept (Gibco), 10µg/ml Fungizone (Gibco), 5% FBS (Gibco)", up to DIV9, when cells were processed for analysis. When due, medium was further supplemented from DIV4 to DIV9 by 10 µM Lamivudine (L1295-10MG, Sigma Aldrich), assuming a conventional 3 days drug halflife.

In general, lentiviral transgenes were activated at day in vivo 0 (DIV0, i.e. the dissection day) by 2µg/mL doxycyclin (Sigma #D9891-10G) medium supplementation, and kept on by further doxycyclin supplementation, performed assuming a conventional 2 days drug halflife.

### Primary neocortical cultures: neuron-enriched ones

Cortical tissue from E16.5 mice was chopped to small pieces for 5 min, in the smallest volume of ice-cold "1X PBS - 0,6% D-glucose - 5mg/ml DNaseI (Roche #10104159001)". After enzymatic digestion in "2.5X trypsin (Gibco #15400054) - 2mg/ml DNaseI" for 5 min, and its inhibition with "DMEM-glutaMAX (Gibco) – 10% FBS (Euroclone) - 1X Pen-Strep", cells were spun down and transferred to differentiative medium [Neurobasal-A, 1X Glutamax (Gibco), 1X B27 supplement (Invitrogen), 25µM L-glutamate (Sigma), 25µM β-Mercaptoethanol (Gibco), 2% FBS, 1X Pen/Strept (Gibco), 10µg/ml fungizone (Gibco)]. Cells were counted and plated onto 0.1mg/ml poly-L-Lysine (Sigma #P2636) pre-treated 12-multiwell plates (Falcon), 8×10^5^ cells per well in 0.6-0.8 ml differentiative medium. 10µM Cytosine β-D- arabinofuranoside (AraC; Sigma #C6645) was added to the medium at DIV1. Cells were kept in culture 8 days.

When required, lentiviral culture transduction was performed at DIV1, and TetON-regulated transgenes were activated by 2µg/ml doxycyclin (Clontech #63131l) medium supplementation at DIV4.

### Lentiviral vectors

Third generation self-inactivating (SIN) lentiviral vectors (LVs) were generated as previously described [79] with minor modifications. Resuspended in "DMEM glutaMAX - 10x FBS - 1x Pen/Strep" HEK293T cells were plated on 10cmØ plates, 8*10^6^ cells/plate. Three days later, they were co-transfected with the transfer vector plasmid plus three auxiliary plasmids (pMD2-VSV.G; pMDLg/pRRE; pRSV-REV), in the presence of LipoD293^TM^ (SigmaGen #SL100668). The conditioned medium was collected 24 and 48 hours after transfection, filtered and ultracentrifuged at 50.000 g on a fixed angle rotor (JA25.50 Beckmann Coulter) for 150 min at 4°C. Lentiviral pellets were then resuspended in (BSA-free) 1X PBS (Gibco). LVs were titrated by Real Time quantitative PCR after infection of HEK293T cells, as previously reported [80]. One end point fluorescence titrated LV was included in each PCR titration session and PCR-titers were adjusted to fluorescence-equivalent titers throughout the study. The full list of LVs employed for this study is reported in **Table 2**. Performances of *Foxg1*-modulating LV transgenes were monitored by qRT-PCR. Results are summarized in **Table S1**.

**Table 2.**
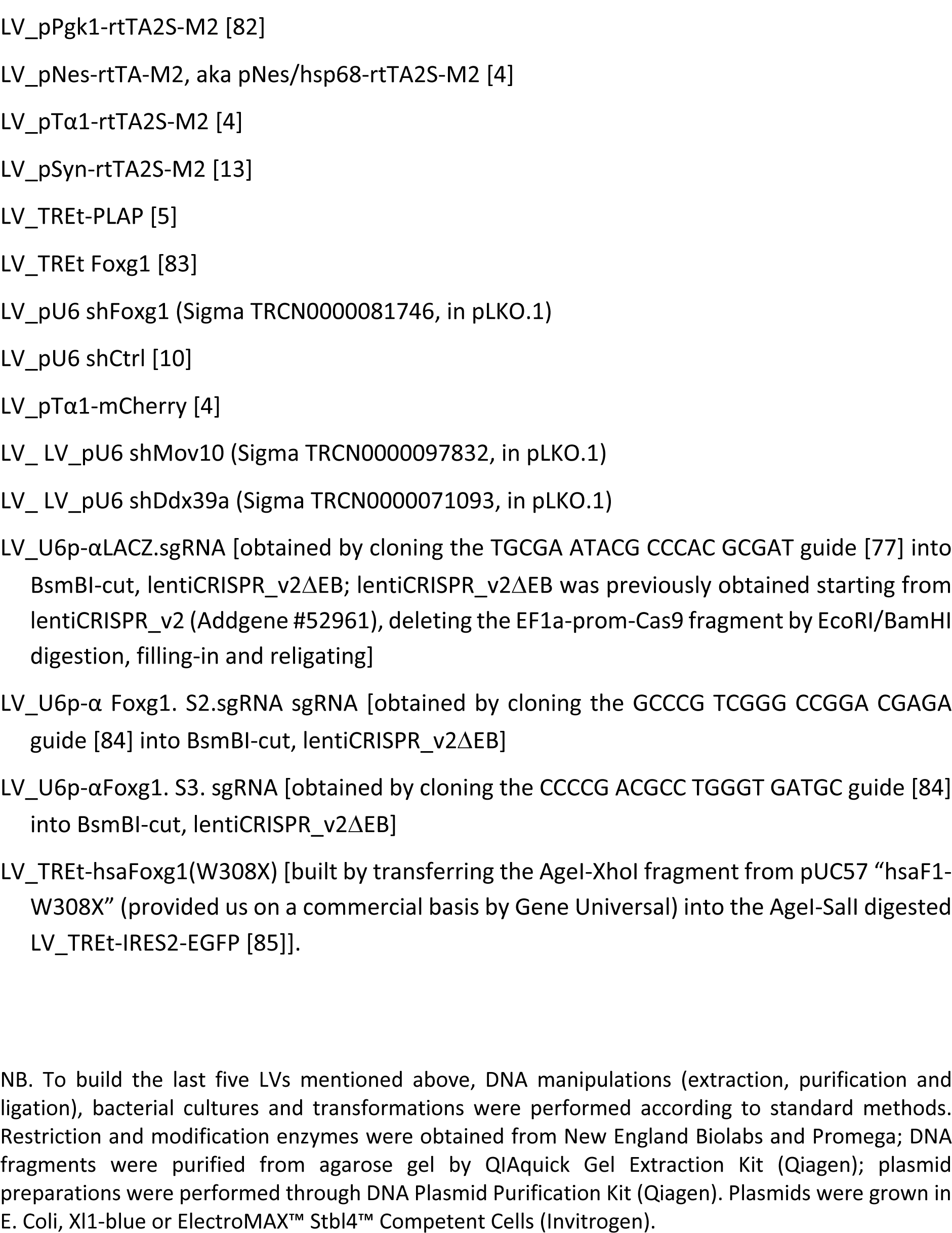
Lentiviruses list. The lentiviruses (LVs) were used for this study were named according to the standard nomenclature: LV: pX-GOI, where pX is the promoter and GOI is the gene of interest. They were:

### Genomic DNA isolation and qPCR amplicon quantitation

DNA was isolated from neocortical tissue as well as from primary pallial cultures. Neocortices from E14.5 embryos and P0 pups were microdissected, cut into small pieces for <5 min in the smallest volume of ice-cold 1X PBS - 0.6% glucose, and kept on ice. Minced neural tissue was further dissociated to single cells by 2X trypsin at 37°C for 5 min, followed by gentle pipetting and enzyme inactivation by FBS. On the other side, primary cell cultures were straightly treated by 0.3X trypsin at 37°C for 5 min, again followed by gentle pipetting and enzyme inactivation by FBS. Next, in both cases, cells were counted in a Burker-chamber, split into aliquots of 10^6^ cells each, pelletted for 5 min at 200g, and stored at -80°C for subsequent use. Single cell aliquots were processed by the “FlexiGene DNA Kit; Qiagen”, according to Manufacturer’s instructions, with the following modifications:

1. PK was employed at 0.6mg/mL (high PK) and 1.2 mg/mL (very high PK);
2. PK sample incubation was extended to 6 hrs;
3. following precipitation, the DNA pellet was washed three times by 70% ethanol.

Finally, DNA was resuspended in water and quantified by DS-11spectrophotometer (DeNovix).

Quantification of genomic amplicons (*L1* elements and X-chromosome genes) was performed starting from 10ng DNA per each reaction, by means of the SsoAdvanced SYBR Green SupermixT“platform (Biorad), according to Manufacturer’s instructions. Primer sequences are reported in **Table 1**. Each amplicon was qPCR-analyzed in technical triplicate, and results averaged. Averages were normalized against levels of selected autosomal amplicons, as reported in Figures (depending on cases, the level of a single reference gene or the geometrical average of >1 of them were employed). As specified in Legends to Figures, biological replicates were DNA preparations originating (1) from different embryo/neonate individuals or (2) individually transduced and cultured cell aliquots, taken from pooled neural cells preparations.

### Chromatin Immunoprecipitation (ChIP) assay

Chromatin Immunoprecipitation-quantitative Polymerase Chain Reaction assays (ChIP-qPCRs) were performed on chromatin aliquots prepared from 3.0*10^5^ cells (αFoxg1-ChIP) or 1.0*10^5^ cells (αH3K4me3-, αH3K9me3-, αH3K27ac-, and αMeCP2-ChIP).

ChIP analysis was performed by the MAGnify^TM^ Chromatin Immunoprecipitation System kit (Invitrogen), according to Manufacturer’s instructions, with minor modifications. Briefly, chromatin was fixed by 1% formaldehyde for 10 min at RT. After cell lysis, fixed chromatin was sonicated by a Soniprep 150 apparatus according to the following settings: (1) on ice; 5s ON, 55 s OFF; oscillation amplitude 5 μm; 4 cycles (αFoxg1-ChIP); (2) on ice; 5s ON, 55 s OFF; oscillation amplitude 5 μm; 5 cycles (αH3K4me3-, αH3K9me3-, αH3K27ac-, and αMeCP2-ChIP). Agarose gel electrophoresis was employed to estimate quality of sonicated chromatin. Sonicated chromatin was immunoprecipitated for 2h at 4°C in a final volume of 100 μL, keeping the tubes in a rotating device, using the following, agarose bead-bound antibodies:

- αFoxg1 (rabbit polyclonal, Abcam #ab18259), 10 μg/reaction;
- αH3K4me3 (rabbit polyclonal, Abcam #ab8580), 3 μg/reaction;
- αH3K9me3 (rabbit polyclonal, Active Motif #39161), 3 μg/reaction;
- αH3K27ac (rabbit polyclonal, #ab177178), Abcam, 3 μg/reaction;
- αMecP2 (rat polyclonal IgG2a serotype, Active Motif #61291), 3 μg/reaction.

Next, immunoprecipitated DNA was purified according to Manufacturer’s instructions. Last, 1/30 of each immunoprecipitated (IP) DNA sample was amplified by qPCR. For each sample, qPCRs were performed in technical triplicate. Averages were normalized against input chromatin and further normalized against controls. Experiments were performed at least in biological triplicate. Results were evaluated by Student’s t-test, via Excel software.

### Total RNA extraction

Total RNA was extracted from both primary neural cultures and acutely dissected neocortical samples using TRIzol Reagent (Thermofisher #15596026) according to the Manufacturer’s instructions. RNA was precipitated using isopropanol and GlycoBlue (Ambion) overnight at - 80°C. After two washes with 75% ethanol, RNA was resuspended in 20µl sterile nuclease-free deionized water. Agarose gel electrophoresis and spectro-photometric measurements (DS-11, DeNovix) were employed to estimate its concentration, quality and purity.

### RNA Immunoprecipitation (RIP)

RNA immunoprecipitation was performed starting from primary neural cultures.

Before starting cells processing, for each RIP reaction, 10µl of protein A/G Dynabeads (Thermofisher #492024) were coupled with 10µg of αFoxg1 (ChIP-grade, rabbit polyclonal, Abcam #ab18259), or 10µg of rabbit IgG (Millipore #12370) as control, according to Manufacturer’s protocols. "Pre-clearing" control beads were prepared omitting antibody coupling.

Cells were washed once with ice-cold 1x PBS. 75µl ice-cold lysis buffer was added to each well (of 12-multiwell plate) and kept on ice for 10 min. Next, cells were scraped and lysed by vigorously pipetting up and down, paying attention not to make bubbles. Lysate collected from 10 wells (about 8×10^6^ cells; corresponding to a αFoxg1/IgG biological samples pair), was “pipetted up and down and kept 10 min on ice" twice, then it was centrifuged at 2000g for 10 min at 4°C, and then centrifuged at 16000g for 10 min at 4°C. The supernatant resulting from each sample was incubated with pre-clearing beads (pre-equilibrated in lysis buffer) for 30 min at 4°C on roller-shaker. Then, preclearing beads were separated with a magnet, and supernatant was incubated with antibody-coupled beads (pre-equilibrated in lysis buffer), overnight at 4°C on roller-shaker. 10% of supernatant (Input, IN-RIP) was stored on ice. The day after, beads were collected with a magnet and washed five times with 0.5ml ice-cold high-salt buffer. [Lysis buffer: 25mM TRIS-HC1, 150mM KCl (Ambion), 10mM MgCl_2_ (Ambion), 1% (vol/vol) NP-40 (Thermo Fisher Scientific), 1X EDTA-free protease inhibitors (Roche), 0.5 mM DTT (Invitrogen), 10 μl/ml rRNasin (Promega), 10 µl/ml SuperaseIn (Applied Biosystems). High-salt buffer: 25mM TRIS-HCI, 350mM KCl (Ambion), 10mM MgCl_2_ (Ambion), 1% (vol/vol) NP-40 (Thermo Fisher Scientific), 1X EDTA-free protease inhibitors (Roche), 0.5mM DTT (Invitrogen)]. For each sample, RNA immunoprecipitated (IP-RIP) and Input were extracted with Trizol^TM^ LS reagent according to manufacturer’s instructions. For each sample, a supplementary extraction was used to improve the total RNA yield. RNA was precipitated using isopropanol and GlycoBlue over-night at -80°C. After two washes with 75% ethanol, the RNA was resuspended in 10µl sterile nuclease-free deionized water. Agarose gel electrophoresis and spectrophotometric measurements (NanoDrop ND-1000) were employed to estimate quantity, quality and purity of the resulting RNA.

### Total and immunoprecipitated RNA quantitation

RNA preparations from total RNA samples, and RIP samples were treated by TURBO^TM^DNase (2U/µl) (Thermofisher # AM2238) for 1h at 37°C, following Manufacturer’s instructions. cDNA was produced by reverse transcription (RT) by Superscript III^TM^ (Thermofisher #18080093) according to Manufacturer’s instructions, in the presence of random hexamers. Then, the RT reaction was diluted 1:5 (in case of both RIP and total RNA samples), and 1µl of the resulting cDNA was used as substrate of any subsequent quantitative PCR (qPCR) reaction. Limited to intron-less amplicons and for RIP-derived samples, negative control PCRs were run on RT(-) RNA preparations. qPCR reactions were performed by the SsoAdvanced SYBR Green Supermix^TM^ platform (Biorad #1725270), according to Manufacturer’s instructions. For each transcript under examination and each sample, cDNA was qPCR-analyzed in technical triplicate, and results averaged. In case of total RNA, mRNA levels were normalized against the geometrical average of *Rpl10a*,*Gapdh*, *Cltc* and *Rn18S* levels (or a subset of them, see Legends to Figures). In case of RIP samples, IP samples were normalized against Inputs. Primer sequences reported in **Table 1**. Data analysis was performed using Microsoft Excel.

### Immunofluorescence: sample preparation and analysis

Neural cells were dissociated by 0.3X trypsin digestion for 4 minutes at RT, followed by gentle (10 times) pipetting and 1:1 v/v trypsin inactivation by FBS-containing medium. Cells were resuspended at 200 cells/μL and 1 mL of suspension was plated on a 12mmØ glass coverslip previously coated by 0.1mg/ml poly-L-Lysine. Cells were kept in 5% CO2, at 37°C for 1 hour, fixed in 4% PFA at RT for 15 minutes, and finally washed 3 times in 1x PBS.

Fixed/washed cells were treated with blocking mix (1X PBS - 10% FBS -1 mg/ml BSA - 0.1% Triton X-100) for at least 1 hour at room temperature (RT). Next, they were incubated with primary antibodies in blocking mix, overnight at 4°C. The day after, samples were washed 3 times in "1X PBS – 0.1% Triton X-100" (5 min each) and then incubated with secondary antibodies in blocking mix, for 2 hours at RT. Samples were finally washed 3 times in "1X PBS – 0.1% Triton X-100" (5 min each), subsequently counterstained with DAPI (4’, 6’-diamidino-2-phenylindole), and mounted in Vectashield Mounting Medium (Vector). The following primary antibodies were used: αSox2, rabbit polyclonal, (clone 2Q178, Abcam #ab6142, 1:400); αTubb3, mouse monoclonal, (clone Tuj1, Covance #MMS-435P, 1:1000); αFoxg1 antibody (rabbit polyclonal, Abcam #ab18259, 1:500); αRFP (specifically recognizing mCherry), rat monoclonal (Antibodies online #ABIN334653, 1:500); αEGFP (Enhanced Green Fluorescent Protein), chicken polyclonal (GenTex #GTX13970, 1:1000). Secondary antibodies were conjugates with Alexa Fluor 488, Alexa Fluor 594 and Alexa Fluor 647 fluorophores (Invitrogen, 1:500).

### Image acquisition and analysis

Immuno-stained cells were photographed on a Nikon Eclipse TI microscope, equipped with a 20X objective, by a Hamamatsu 1394 ORCA-285 camera (**Fig. 2**), or a Nikon C1 confocal system (**Fig. 5**). Hamamatsu photos were collected as 1344×1024 pixel files. Nikon C1 photos were collected as 3μm Z-stacks (0.3μm steps) of 1024×1024 pixel images. Images were analyzed by Adobe Photoshop CS6 (**Fig. 2**) and Volocity 5.5.1 (**Figure 5**) softwares. Resulting numerical data were further processed by Microsoft Excel software.

Specifically in case of **Fig. 5** analysis, the following strategy was implemented. Acutely transduced by LV_pTα1-mCherry (firing in NPS and Ns) at moi=8, NSCs, NPs and Ns were recognized by their mCherry^−^Tubb3^−^, mCherry^+^Tubb3^−^ and mCherry^±^Tubb3^+^ profiles, respectively. Moreover, for each cell, nuclear Foxg1 protein content was quantified by Volocity 5.5.1 analysis of aFoxg1-IF signal. Data referring to an equal number of cells from 6 biological replicates of control samples were collected, cumulatively ranked and employed to establish boundaries between contiguous (aFoxg1-IF signal) deciles (here "biological replicates" are aliquots of neural cells originating from the same starting pool, each independently transduced and cultured). Next, starting from 6 and 5 biological replicates of control and *Foxg1*-OE samples, respectively, distinctive cell types (NSCs, NP, and Ns) of different genotypes (control or mis-expressing Foxg1), falling within different decile bins were quantified, normalized against total cells of the same type and genotype, and finally plotted against decile number. Cumulatively, >17,000 neural cells were scored for this analysis.

### 3.17. Statistical analysis

When not otherwise stated, experiments were performed at least in biological triplicate. Statistical tests employed for result evaluation, *p* values, and definitions of *n* (number of biological replicates) are provided in each Figure. Full primary data referred to by all Figures are reported in **Table S4**.

## AUTHORS’ CONTRIBUTIONS

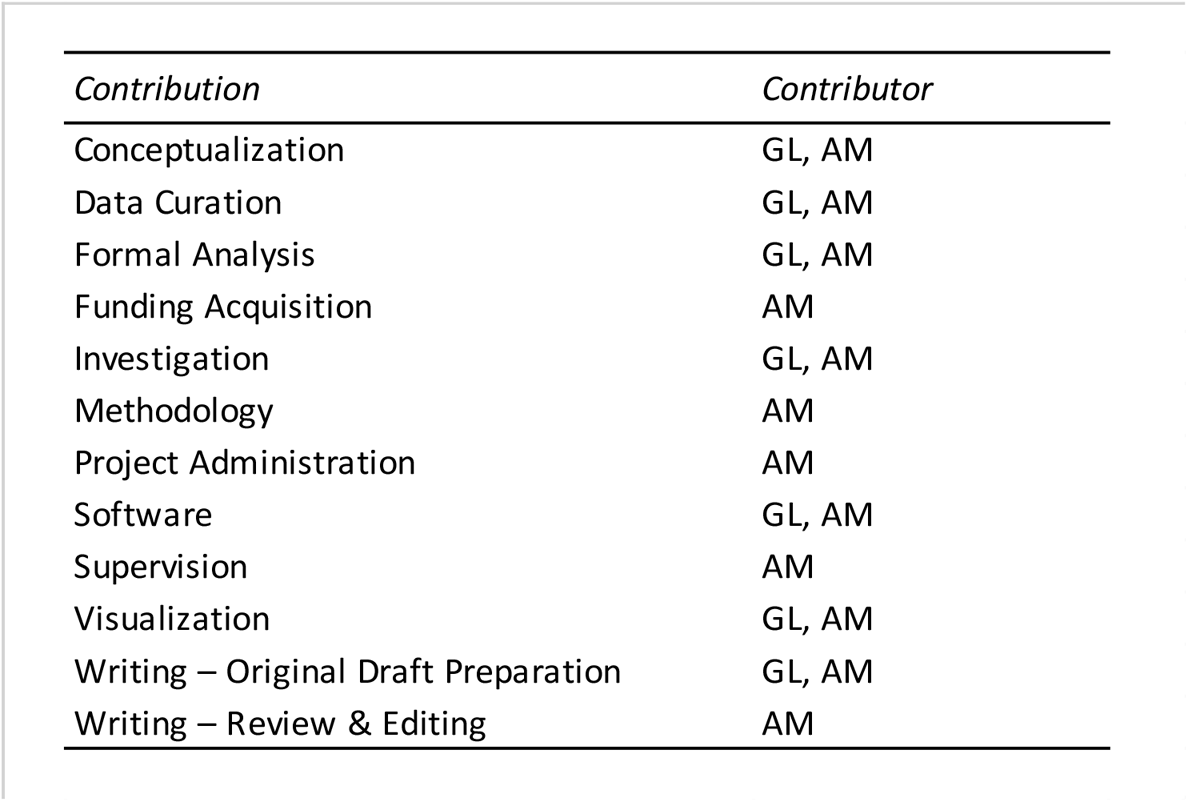

## ACKNOWLEDGEMENTS

We thank Osvaldo Artimagnella for methodological help with RIP assays and astroglia-free neuronal cultures, and Laura Rigoldi for embryo sexing.

## FUNDING

We thank:

1. International FOXG1 Research Foundation (Grant to A.M.)
2. SISSA (intramurary funding to A.M.)

## CONFLICTING INTERESTS

The Authors declare no conflict of interests.

## Notes

### Competing Interest Statement

The authors have declared no competing interest.

