## Supplementary material for "*Foxg1* bimodally tunes *L1*-mRNA and -DNA dynamics in the developing murine neocortex": 2 supplemental figures & 4 supplemental tables

**including:**

**- 2 Supplementary Figures, with Legends**

**- 4 Supplementary Tables, with Legends**

**SUPPLEMENTARY FIGURES**

**Figure S1**

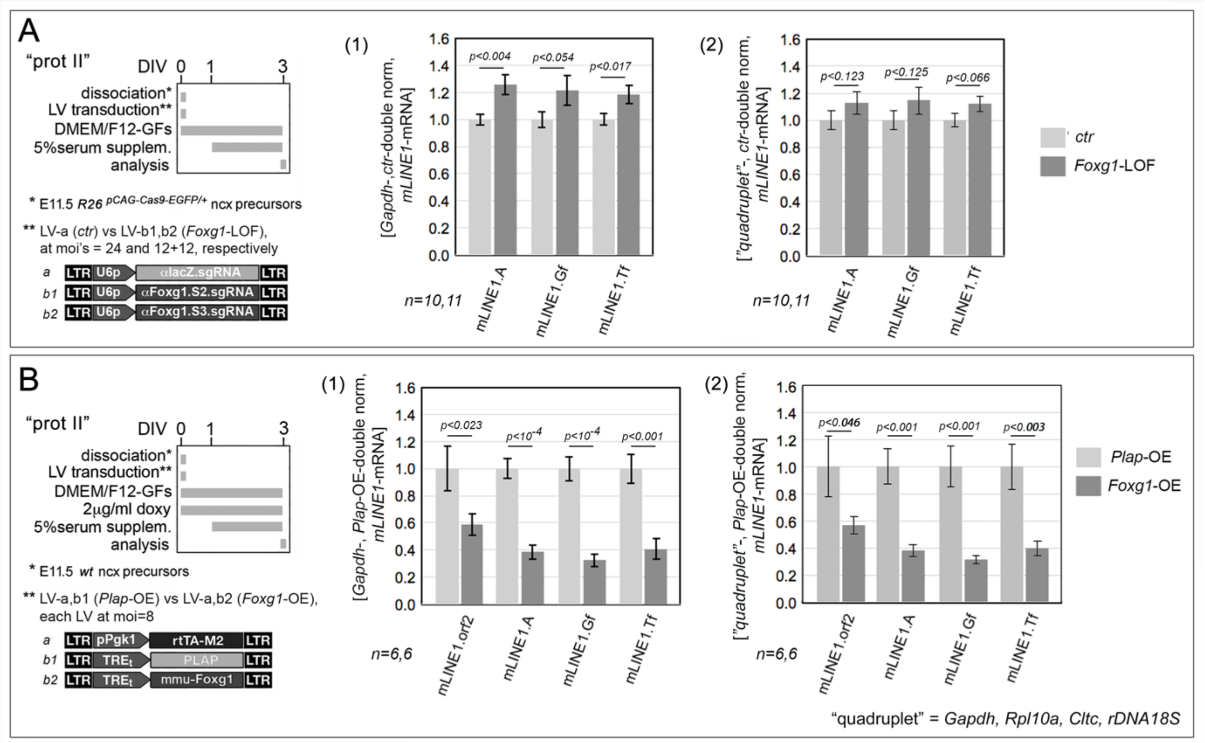

**Legend to Figure S1. Comparative evaluation of "quadruplet-" vs *Gapdh*-normalization of *L1*-mRNA expression levels.** In (A) and (B) shown are data referred to in Fig. 3A and 3D, respectively, upon alternative normalization against *Gapdh* (1) and gene "quadruplet" (2).

**Figure S2**

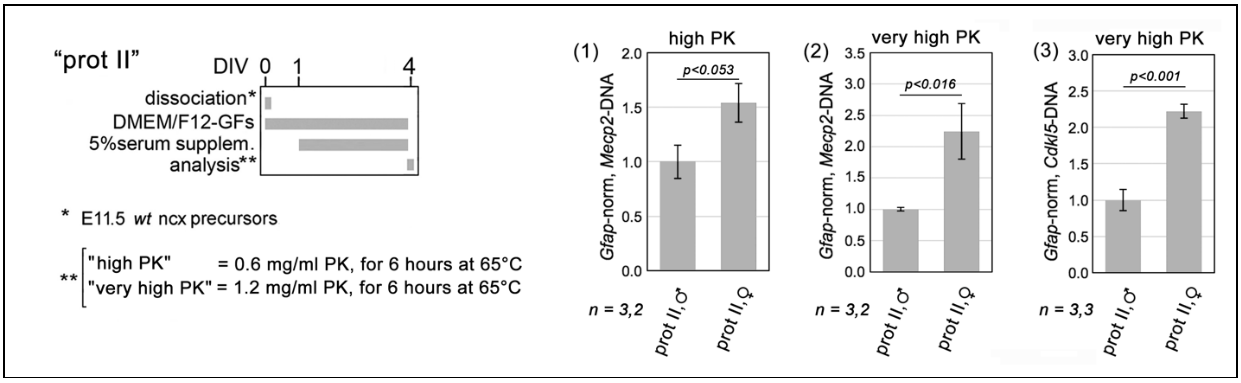

**Legend to Figure S2. Optimization of genomic DNA extraction.** To left, protocol. Mid-neuronogenic cultures set by means of type II protocols. DNA extraction performed by a strong ("high PK", see Materials and Methods) or a very strong procedure ("very high PK", see Materials and Methods). To right, results. Graphs (1-3) providing a comparative assessment of "high PK" and "very high PK" procedures for their capability to extract with similar efficacy genomic DNA, regardless of chromatin accessibility. Such assesment was done by quantifying X-chromosomal *Mecp2* and *Cdkl5* sequences, amplified from female and male genomic DNA, and normalizing them against the euchromatic *Gfap* autosomal locus. Error-bars representing sem's. Statistical significance of results evaluated by t-test (one-tailed, unpaired). *n* is the number of biological replicates, i.e. independently cultured and engineered cell aliquots, originating from pooled, *wild-type* E11.5 neocortical primordia.

#### SUPPLEMENTARY TABLES

**Table S1.**

| *ref* | | *Foxg1-mRNA manipulation* | |
|---|---|---|---|
| *Fig. #* | *panel, graph* | *ctr-norm Foxg1-mRNA levels* | *normalizer(s)* |
| 1 | --- | 0.64 | *Gapdh* |
| 3 | A | 0.64 | *gene quadruplet** |
| | B | 2.74 | *gene quadruplet** |
| | C | 3.63 | *gene quadruplet** |
| | D | 2.84 | *gene quadruplet** |
| | E | 4.02 | *gene quadruplet** |
| 4 | graph (1) | 4.84 | *Gapdh* |
| | graph (2) | 4.27 | *Rpl10a* |
| | graph (3) | 0.28 | *Rpl10a* |
| 9 | --- | 14.08 | *gene quadruplet** |
| 12 | --- | 0.64 | *Gapdh* |
| 13 | B, graph (1) | 0.35 | *Gapdh* |
| | B, graph (3) | 3.05 | *Gapdh* |

**(Gapdh; Rpl10a; Cltc and rDNA 18S)*

**Legend to Table S1. Foxg1-mRNA dynamics upon OE- and LOF-manipulations of this study.** Here, reported are control-normalized *Foxg1*-mRNA levels in selected experiments of this study, each with reference Figures and panels, and normalizer gene(s) employed for their evaluation.

**Table S2.**

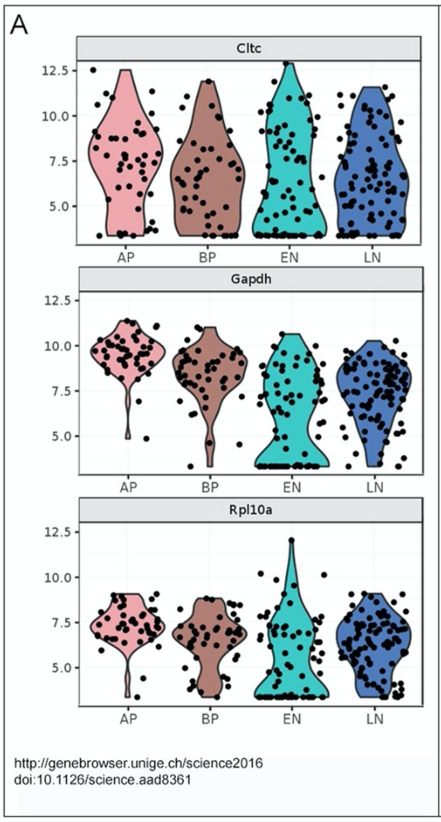

B

| Locus | log2FC | logCPM | P-value | FDR | Gene Name |
|---|---|---|---|---|---|
| ENSMUSG00000047126 | 0,01 | 9,73 | 0,94 | 0,96 | Cltc |

| Locus | log2FC | logCPM | P-value | FDR | Gene Name |
|---|---|---|---|---|---|
| ENSMUSG00000057666 | -0,16 | 9,44 | 0,15 | 0,24 | Gapdh |

| Locus | log2FC | logCPM | P-value | FDR | Gene Name |
|---|---|---|---|---|---|
| ENSMUSG00000037805 | 0,07 | 6,50 | 0,68 | 0,77 | Rpl10a |

doi:10.5281/zenodo.3739467

**Legend to Table S2. "Quadruplet", RNA-pol II-transcribed normalizers: fluctuations of their mRNA levels throughout normal neuronogenesis as well as upon *Foxg1* manipulation in differentiating neurons. (A)** Violin plots representing single cell mRNA levels, in pallial apical precursors (AP), basal precursors (BP), early neurons (EN) and late neurons (LN). Downloaded from http://genebrowser.unige.ch/science 2016 (Telley et al., 2016). **(B)** Modulation of average mRNA levels in differentiating neuronal cultures overexpressing *Foxg1* (Artimagnella and Mallamaci, 2020 i.e doi:10.5281/zenodo.3739467).

**Table S3**

| *Foxg1-responsive gene (ref 1)* | *impact on retro-transcription/transposition* | *mRNA dynamics upon Foxg1 overexpression in ncx neurons* | | | | |
|---|---|---|---|---|---|---|
| | | *Locus* | *log2FC* | *logCPM* | *P-value* | *FDR* |
| Apobec1 | limits Line1 retrotransposition (ref 2) | ENSMUSG00000040613 | -1.01 | 2.02 | 0.00 | 0.00 |

| *Foxg1 interactor (ref 3)* | *impact on retro-transcription/transposition* | *Foxg1-protein interaction details (Biogrid #)* |
|---|---|---|
| MOV10 | limits Line1 retrotransposition (ref 4) | 1507467 |
| DDX39A | limits Line1 retrotransposition (ref 4) | 1507432 |

*REFERENCES*

1 doi: 10.5281/zenodo.2849656.
2 doi: 10.1093/nar/gkr124
3 https://thebiogrid.org/
4 doi: 10.1371/journal.pgen.1002941

**Legend to Table S3. Putative mediators of Foxg1 impact on L1 retro-transcription.** Shown are selected genes impacting on L1 retro-transcription/transposition rates, (A) whose mRNA levels are sensitive to Foxg1 overexpression, or (B) whose protein products physically interact with Foxg1 protein.

#### Table S4. Full primary data (ref: Figures 1,2,3,4,5,6,8,9,10,11,12,13,14,15,S1,S2)

*Figure 1*

P0 neocortex

Gadph-norm, Ctr-norm [mRNA]

| mLine1.orf2 Foxg1 (+/+) | mLine1.orf2 Foxg1 (+/-) | mLine1.A Foxg1 (+/+) | mLine1.A Foxg1 (+/-) | mLine1.Gf Foxg1 (+/+) | mLine1.Gf Foxg1 (+/-) | mLine1.Tf Foxg1 (+/+) | mLine1.Tf Foxg1 (+/-) |
|---|---|---|---|---|---|---|---|
| 1.00 | | | | | | | |
| 0.87 | 1.27 | | 1.28 | 1.17 | 1.45 | 1.07 | 1.44 |
| 0.81 | 1.05 | 0.99 | 0.94 | 0.97 | 1.13 | 0.98 | 1.03 |
| 1.10 | 0.97 | 0.94 | 1.03 | 0.87 | 1.29 | 0.83 | 1.05 |
| 0.82 | 1.24 | 0.97 | 1.17 | 1.07 | 1.32 | 1.07 | 1.26 |
| 0.95 | 1.21 | 1.01 | 1.22 | 0.85 | 1.35 | 0.88 | 1.10 |
| 1.27 | 1.36 | 0.98 | 1.25 | 0.99 | 1.34 | 1.05 | 1.24 |
| 1.17 | 1.28 | 1.11 | 1.31 | 1.07 | 1.43 | 1.13 | 1.34 |

*Figure 2*

**A**

"prot I"

cell freq

| Sox2+ Tubb3- | Sox2+ Tubb3+ | Sox2- Tubb3+ | Sox2- Tubb3- |
|---|---|---|---|
| 0.41 | 0.28 | 0.02 | 0.29 |
| 0.38 | 0.31 | 0.02 | 0.29 |
| 0.39 | 0.27 | 0.02 | 0.32 |

"prot II"

cell freq

| Sox2+ Tubb3- | Sox2+ Tubb3+ | Sox2- Tubb3+ | Sox2- Tubb3- |
|---|---|---|---|
| 0.23 | 0.26 | 0.14 | 0.39 |
| 0.24 | 0.27 | 0.15 | 0.33 |
| 0.19 | 0.27 | 0.16 | 0.41 |

"prot III"

cell freq

| Tubb3+ | Tubb3- |
|---|---|
| 0.72 | 0.28 |
| 0.72 | 0.28 |
| 0.72 | 0.28 |

**B**

(1)

Gapdh-, "prot I"-double-norm mRNA

| orf2 E12 | orf2 E14 | orf2 E20 |
|---|---|---|
| 0.95 | 20.59 | 18.91 |
| 0.83 | 24.61 | 26.32 |
| 1.07 | 18.40 | 30.29 |
| 0.67 | 25.92 | 27.82 |
| 1.48 | 8.18 | 17.66 |
| 1.00 | 21.40 | 8.92 |

(2)

Gapdh-, "prot I"-double-norm mRNA

| mLine1.A-5'UTR E12 | mLine1.A-5'UTR E14 | mLine1.A-5'UTR E20 | mLine1.Gf-5'UTR E12 | mLine1.Gf-5'UTR E14 | mLine1.Gf-5'UTR E20 | mLine1.Tf-5'UTR E12 | mLine1.Tf-5'UTR E14 | mLine1.Tf-5'UTR E20 |
|---|---|---|---|---|---|---|---|---|
| 0.54 | 10.40 | 36.28 | | 7.51 | 23.39 | | 15.66 | |
| 2.95 | 10.18 | 44.18 | 1.11 | 6.91 | 28.72 | 1.02 | 13.70 | 52.68 |
| 0.63 | 9.36 | 32.65 | 0.96 | 7.23 | 16.95 | 1.04 | 14.26 | 78.68 |
| 0.57 | 13.33 | 41.51 | 1.05 | 9.79 | 23.11 | 1.04 | 19.29 | 56.39 |
| 0.81 | 17.40 | 89.81 | 1.25 | 26.47 | 63.74 | 1.11 | 59.45 | 59.86 |
| 0.50 | 17.39 | 54.80 | 0.64 | 9.03 | 43.93 | 0.79 | 18.11 | 114.81 |

**C**

neocortex

quadruplet-norm, ctrl-norm [mRNA]

| mLine1.orf2 E14 | mLine1.orf2 P0 | mLine1.A E14 | mLine1.A P0 | mLine1.Gf E14 | mLine1.Gf P0 | mLine1.Tf E14 | mLine1.Tf P0 |
|---|---|---|---|---|---|---|---|
| 1.16 | | 1.01 | | | | 1.18 | |
| 1.05 | | 0.95 | | 1.24 | | 1.11 | |
| 0.98 | | 0.95 | | 0.81 | | 0.93 | |
| 1.10 | 3.03 | 1.13 | 3.16 | 0.97 | 3.14 | 1.05 | 2.53 |
| 0.77 | 1.48 | 0.78 | 1.86 | 1.19 | 1.89 | 0.85 | 1.53 |
| 0.91 | 1.44 | 0.92 | 1.92 | 0.77 | 1.43 | 0.92 | 1.43 |
| 1.17 | 1.77 | 1.30 | 2.34 | 0.86 | 2.19 | 1.05 | 1.77 |
| 0.88 | 4.78 | 1.06 | 4.68 | 1.09 | 5.29 | 0.98 | 3.32 |
| 0.84 | 1.65 | 0.84 | 1.32 | 0.90 | 2.09 | 1.02 | 1.47 |
| 1.13 | 1.41 | 1.06 | 1.57 | 1.17 | 1.44 | 0.93 | 1.32 |

mesencephalon

quadruplet-norm, ctrl-norm [mRNA]

| mLine1.orf2 E14 | mLine1.orf2 P0 | mLine1.A E14 | mLine1.A P0 | mLine1.Gf E14 | mLine1.Gf P0 | mLine1.Tf E14 | mLine1.Tf P0 |
|---|---|---|---|---|---|---|---|
| | | | | 0.89 | | 0.95 | |
| 0.63 | | 0.77 | | 0.81 | | 0.81 | |
| 0.76 | 0.64 | 0.81 | 0.82 | 1.33 | 0.75 | 1.24 | 0.81 |
| 1.64 | 0.67 | 1.34 | 0.72 | 0.95 | 0.56 | 1.52 | 0.66 |
| 0.84 | 1.20 | 0.84 | 0.98 | 0.38 | 0.76 | 0.37 | 0.86 |
| 0.29 | 0.71 | 1.44 | 0.71 | 1.82 | 0.54 | 1.46 | 0.67 |
| 1.74 | 0.95 | 1.19 | 0.78 | 1.28 | 0.58 | 1.10 | 0.95 |
| 1.40 | 0.73 | 1.14 | 0.58 | 1.00 | 0.45 | 1.08 | 0.52 |
| 0.96 | 1.19 | 0.92 | 1.02 | 0.96 | 0.83 | 0.95 | 0.92 |
| 0.73 | 0.71 | 0.55 | 0.73 | 0.58 | 0.60 | 0.52 | 0.67 |

*Figure 3*

**A**

["quadruplet"-, ctrl-double-norm *mLINE1*-mRNA]

| mLine1.A ctr | mLine1.A Foxg1-LOF | mLine1.Gf ctr | mLine1.Gf Foxg1-LOF | mLine1.Tf ctr | mLine1.Tf Foxg1-LOF |
|---|---|---|---|---|---|
| | 1.18 | | 1.22 | | 1.11 |
| 0.77 | 1.35 | 1.11 | 1.41 | 0.89 | 1.37 |
| 0.89 | 1.43 | 0.84 | 1.00 | 0.99 | 1.05 |
| 0.97 | 1.58 | 1.07 | 1.37 | 1.03 | 1.38 |
| 0.83 | 0.86 | 0.73 | 1.79 | 0.93 | 1.34 |
| 1.31 | 0.77 | 1.12 | 0.60 | 1.01 | 0.81 |
| 0.83 | 1.09 | 0.80 | 1.17 | 0.90 | 1.12 |
| 0.82 | 0.82 | 0.76 | 0.80 | 0.80 | 0.88 |
| 1.02 | 1.31 | 0.97 | 1.31 | 0.93 | 1.11 |
| 1.32 | 1.09 | 1.27 | 1.11 | 1.26 | 1.20 |
| 1.23 | 0.93 | 1.34 | 0.84 | 1.26 | 0.96 |

**B**

["quadruplet"-, *Plap*-OE-double norm mLINE1- mRNA]

| mLine1.orf2 ctr | mLine1.orf2 Foxg1-OE | mLine1.A ctr | mLine1.A Foxg1-OE | mLine1.Gf ctr | mLine1.Gf Foxg1-OE | mLine1.Tf ctr | mLine1.Tf Foxg1-OE |
|---|---|---|---|---|---|---|---|
| 1.05 | 0.89 | 1.00 | 0.95 | 0.97 | 1.04 | 0.98 | 1.08 |
| 0.92 | 0.75 | 1.00 | 0.68 | 0.99 | 0.75 | 0.96 | 0.69 |
| 0.94 | 0.75 | 0.81 | 0.69 | 0.83 | 0.70 | 0.73 | 0.72 |
| 0.89 | 0.64 | 0.97 | 0.69 | 0.97 | 0.74 | 0.98 | 0.77 |
| 1.08 | 0.87 | 1.32 | 0.87 | 1.47 | 0.98 | 1.37 | 1.00 |
| 1.18 | 0.71 | 1.03 | 0.72 | 0.96 | 0.70 | 1.08 | 0.74 |
| 0.93 | 0.59 | 0.93 | 0.62 | 1.00 | 0.55 | 0.96 | 0.65 |
| 1.08 | 0.66 | 1.08 | 0.74 | 1.00 | 0.76 | 1.11 | 0.78 |
| 0.93 | 0.52 | 0.86 | 0.59 | 0.81 | 0.69 | 0.83 | 0.55 |

**D**

["quadruplet"-, *Plap*-OE-double norm mLINE1- mRNA]

| mLine1.orf2 ctr | mLine1.orf2 Foxg1-OE | mLine1.A ctr | mLine1.A Foxg1-OE | mLine1.Gf ctr | mLine1.Gf Foxg1-OE | mLine1.Tf ctr | mLine1.Tf Foxg1-OE |
|---|---|---|---|---|---|---|---|
| 0.95 | 0.74 | 0.87 | 0.31 | 0.99 | 0.28 | 0.89 | 0.38 |
| 0.48 | 0.72 | 0.67 | 0.35 | 0.65 | 0.30 | 0.78 | 0.41 |
| 1.96 | 0.35 | 1.60 | 0.26 | 1.67 | 0.24 | 1.75 | 0.26 |
| 1.04 | 0.57 | 0.98 | 0.48 | 1.08 | 0.44 | 1.14 | 0.65 |
| 1.09 | 0.46 | 1.03 | 0.54 | 0.85 | 0.29 | 0.83 | 0.34 |
| 0.48 | 0.58 | 0.85 | 0.37 | 0.76 | 0.36 | 0.62 | 0.35 |

**C**

["quadruplet"-, *Plap*-OE-double norm mLINE1-mRNA]

| mLine1.orf2 ctr | mLine1.orf2 Foxg1-OE | mLine1.A ctr | mLine1.A Foxg1-OE | mLine1.Gf ctr | mLine1.Gf Foxg1-OE | mLine1.Tf ctr | mLine1.Tf Foxg1-OE |
|---|---|---|---|---|---|---|---|
| 0.90 | 0.75 | 1.01 | 0.65 | 0.88 | 0.68 | 1.05 | 0.74 |
| 1.10 | 0.92 | 1.09 | 0.73 | 1.10 | 0.74 | 1.03 | 0.67 |
| 1.15 | 0.59 | 1.06 | 0.54 | 1.02 | 0.47 | 0.98 | 0.55 |
| 0.97 | 0.77 | 0.89 | 0.99 | 0.87 | 0.81 | 0.97 | 0.85 |
| 0.70 | 0.70 | 0.76 | 0.67 | 0.75 | 0.58 | 0.73 | 0.73 |
| 1.19 | 0.67 | 1.19 | 0.55 | 1.38 | 0.42 | 1.24 | 0.53 |

**E**

["quadruplet"-, *Plap*-OE-double norm mLINE1-mRNA]

| mLine1.orf2 ctr | mLine1.orf2 Foxg1-OE | mLine1.A ctr | mLine1.A Foxg1-OE | mLine1.Gf ctr | mLine1.Gf Foxg1-OE | mLine1.Tf ctr | mLine1.Tf Foxg1-OE |
|---|---|---|---|---|---|---|---|
| 0.98 | | 1.01 | | 0.97 | | 0.99 | |
| 1.05 | | 1.15 | | 1.11 | | 1.03 | |
| 1.21 | 0.85 | 1.15 | 0.76 | 1.09 | 0.74 | 1.15 | 0.80 |
| 0.79 | 1.05 | 0.85 | 0.86 | 0.92 | 0.75 | 0.87 | 0.86 |
| 1.14 | 1.16 | 1.13 | 0.98 | 1.19 | 0.90 | 1.22 | 1.00 |
| 0.83 | 0.79 | 0.74 | 0.72 | 0.78 | 0.69 | 0.76 | 0.75 |
| 1.01 | 0.79 | 0.97 | 0.77 | 0.94 | 0.76 | 0.99 | 0.79 |

**G**

["quadruplet"-, *Plap*-OE-double norm mLINE1- mRNA]

| mLine1.orf2 ctr | mLine1.orf2 Foxg1-OE | mLine1.A ctr | mLine1.A Foxg1-OE | mLine1.Gf ctr | mLine1.Gf Foxg1-OE | mLine1.Tf ctr | mLine1.Tf Foxg1-OE |
|---|---|---|---|---|---|---|---|
| | 0.77 | | 0.87 | | 0.86 | | |
| 0.86 | 0.73 | 0.82 | 0.71 | 0.80 | 0.69 | 0.84 | 0.88 |
| 0.95 | 0.65 | 1.06 | 0.67 | 1.06 | 0.60 | 1.02 | 0.69 |
| 0.88 | 0.71 | 0.86 | 0.69 | 0.87 | 0.72 | 0.86 | 0.63 |
| 1.10 | 0.87 | 1.04 | 0.82 | 1.07 | 0.85 | 1.06 | 0.70 |
| 0.95 | 0.92 | 0.98 | 0.97 | 0.99 | 0.91 | 1.00 | 0.83 |
| 1.02 | 0.88 | 0.97 | 0.88 | 1.00 | 0.84 | 0.96 | 0.89 |
| 1.24 | 0.93 | 1.27 | 1.25 | 1.20 | 1.28 | 1.25 | 0.87 |

**F**

["quadruplet"-, *Plap*-OE-double norm mLINE1- mRNA]

| mLine1.orf2 ctr | mLine1.orf2 Foxg1-OE | mLine1.A ctr | mLine1.A Foxg1-OE | mLine1.Gf ctr | mLine1.Gf Foxg1-OE | mLine1.Tf ctr | mLine1.Tf Foxg1-OE |
|---|---|---|---|---|---|---|---|
| 0.84 | | 0.87 | | 0.89 | 0.87 | 0.83 | |
| 1.05 | 0.86 | 1.10 | 0.89 | 1.07 | 0.89 | 1.05 | 0.91 |
| 0.95 | 0.93 | 1.03 | 0.93 | 1.10 | 0.92 | 1.09 | 1.01 |
| 0.93 | 0.91 | 0.90 | 0.97 | 0.93 | 0.89 | 0.84 | 1.02 |
| 1.02 | 0.90 | 0.94 | 0.97 | 0.89 | 0.82 | 0.93 | 1.05 |
| 1.21 | 0.92 | 1.15 | 0.89 | 1.11 | 0.97 | 1.25 | 0.98 |

*Figure 4*

(1) [*Gapdh*- , ctrl-double-norm *mLINE1.orf2*-mRNA]

| ctr | *Foxg1* -OE |
|---|---|
| 1.03 | 0.65 |
| 0.74 | 0.80 |
| 1.05 | 0.97 |
| 1.18 | 0.57 |

(2) [*Rpl10a*- , ctrl-double-norm *mLINE1.orf2*-mRNA]

| ctr | *Foxg1* -OE |
|---|---|
| 0.97 | 0.54 |
| 0.71 | 0.82 |
| 1.16 | 0.81 |
| 1.16 | 0.49 |

(3) [*Rpl10a*- , ctrl-double-norm *mLINE1.orf2*-mRNA]

| ctr | *Foxg1* -LOF |
|---|---|
| 0.72 | 1.42 |
| 1.10 | 1.47 |
| 1.12 | 2.32 |
| 1.06 | 1.45 |

### *Figure 5*

**A**

(1)

| | NSC | | NP | | N | |
|---|---|---|---|---|---|---|
| | *Plap-OE* | *Foxg1-OE* | *Plap-OE* | *Foxg1-OE* | *Plap-OE* | *Foxg1-OE* |
| frequency | | 0.60 | | 0.38 | | 0.02 |
| | 0.84 | 0.52 | 0.12 | 0.43 | 0.04 | 0.06 |
| | 0.83 | 0.56 | 0.13 | 0.41 | 0.04 | 0.02 |
| | 0.75 | 0.63 | 0.21 | 0.35 | 0.04 | 0.02 |
| | 0.92 | 0.59 | 0.06 | 0.37 | 0.02 | 0.04 |
| | 0.88 | 0.57 | 0.08 | 0.40 | 0.03 | 0.02 |

(2)

| NSC | $^{pNes}$Plap-OE cell n = 6060 | | $^{pNes}$Foxg1-OE cell n = 5811 | | | | | | | |
|---|---|---|---|---|---|---|---|---|---|---|
| cell fraction | $^{pNes}$Plap-OE | 0.12 | 0.12 | 0.12 | 0.12 | 0.12 | 0.11 | 0.10 | 0.09 | 0.08 | 0.03 |
| | $^{pNes}$Foxg1-OE | 0.05 | 0.08 | 0.09 | 0.09 | 0.12 | 0.11 | 0.10 | 0.10 | 0.10 | 0.15 |
| | expr decile | 10 | 9 | 8 | 7 | 6 | 5 | 4 | 3 | 2 | 1 |

| NP | $^{pNes}$Plap-OE cell n = 816 | | $^{pNes}$Foxg1-OE cell n = 3955 | | | | | | | |
|---|---|---|---|---|---|---|---|---|---|---|
| cell fraction | $^{pNes}$Plap-OE | 0.00 | 0.00 | 0.00 | 0.00 | 0.00 | 0.03 | 0.09 | 0.14 | 0.25 | 0.49 |
| | $^{pNes}$Foxg1-OE | 0.00 | 0.00 | 0.00 | 0.00 | 0.00 | 0.01 | 0.02 | 0.05 | 0.09 | 0.83 |
| | expr decile | 10 | 9 | 8 | 7 | 6 | 5 | 4 | 3 | 2 | 1 |

| N | $^{pNes}$Plap-OE cell n = 244 | | $^{pNes}$Foxg1-OE cell n = 296 | | | | | | | |
|---|---|---|---|---|---|---|---|---|---|---|
| cell fraction | $^{pNes}$Plap-OE | 0.01 | 0.01 | 0.02 | 0.03 | 0.04 | 0.05 | 0.07 | 0.14 | 0.18 | 0.46 |
| | $^{pNes}$Foxg1-OE | 0.00 | 0.00 | 0.00 | 0.01 | 0.02 | 0.01 | 0.02 | 0.02 | 0.06 | 0.85 |
| | expr decile | 10 | 9 | 8 | 7 | 6 | 5 | 4 | 3 | 2 | 1 |

**B**

(1)

| | NSC | | NP | | N | |
|---|---|---|---|---|---|---|
| | *Plap-OE* | *Foxg1-OE* | *Plap-OE* | *Foxg1-OE* | *Plap-OE* | *Foxg1-OE* |
| frequency | | 0.12 | | 0.31 | | 0.57 |
| | 0.12 | 0.12 | 0.15 | 0.26 | 0.73 | 0.62 |
| | 0.13 | 0.17 | 0.20 | 0.25 | 0.67 | 0.59 |
| | 0.14 | 0.08 | 0.23 | 0.24 | 0.62 | 0.68 |
| | 0.12 | 0.16 | 0.25 | 0.26 | 0.63 | 0.58 |
| | 0.11 | 0.14 | 0.23 | 0.29 | 0.65 | 0.57 |

(2)

| NSC | $^{pTa1}$Plap-OE cell n = 430 | | $^{pTa1}$Foxg1-OE cell n = 630 | | | | | | | |
|---|---|---|---|---|---|---|---|---|---|---|
| cell fraction | $^{pTa1}$Plap-OE | 0.55 | 0.34 | 0.09 | 0.01 | 0.00 | 0.00 | 0.00 | 0.00 | 0.00 | 0.00 |
| | $^{pTa1}$Foxg1-OE | 0.44 | 0.32 | 0.17 | 0.06 | 0.00 | 0.00 | 0.00 | 0.00 | 0.00 | 0.00 |
| | expr decile | 10 | 9 | 8 | 7 | 6 | 5 | 4 | 3 | 2 | 1 |

| NP | $^{pTa1}$Plap-OE cell n = 745 | | $^{pTa1}$Foxg1-OE cell n = 1286 | | | | | | | |
|---|---|---|---|---|---|---|---|---|---|---|
| cell fraction | $^{pTa1}$Plap-OE | 0.00 | 0.00 | 0.04 | 0.08 | 0.10 | 0.16 | 0.19 | 0.20 | 0.16 | 0.06 |
| | $^{pTa1}$Foxg1-OE | 0.00 | 0.00 | 0.02 | 0.09 | 0.12 | 0.13 | 0.17 | 0.17 | 0.18 | 0.12 |
| | expr decile | 10 | 9 | 8 | 7 | 6 | 5 | 4 | 3 | 2 | 1 |

| N | $^{pTa1}$Plap-OE cell n = 2231 | | $^{pTa1}$Foxg1-OE cell n = 2820 | | | | | | | |
|---|---|---|---|---|---|---|---|---|---|---|
| cell fraction | $^{pTa1}$Plap-OE | 0.05 | 0.08 | 0.12 | 0.12 | 0.12 | 0.10 | 0.09 | 0.08 | 0.10 | 0.13 |
| | $^{pTa1}$Foxg1-OE | 0.03 | 0.08 | 0.08 | 0.10 | 0.10 | 0.10 | 0.09 | 0.07 | 0.11 | 0.24 |
| | expr decile | 10 | 9 | 8 | 7 | 6 | 5 | 4 | 3 | 2 | 1 |

**C**

(1)

| | NSC | | NP | | N | |
|---|---|---|---|---|---|---|
| | *Plap-OE* | *Foxg1-OE* | *Plap-OE* | *Foxg1-OE* | *Plap-OE* | *Foxg1-OE* |
| frequency | | 0.28 | | 0.20 | | 0.52 |
| | | 0.21 | | 0.27 | | 0.52 |
| | | 0.11 | | 0.37 | | 0.52 |
| | 0.10 | 0.09 | 0.27 | 0.29 | 0.62 | 0.61 |
| | 0.10 | 0.14 | 0.27 | 0.31 | 0.63 | 0.54 |
| | 0.12 | 0.17 | 0.28 | 0.32 | 0.60 | 0.51 |

(2)

| NSC | $^{pPgk1}$Plap-OE cell n = 349 | | $^{pPgk1}$Foxg1-OE cell n = 1151 | | | | | | | |
|---|---|---|---|---|---|---|---|---|---|---|
| cell fraction | $^{pPgk1}$Plap-OE | 0.62 | 0.33 | 0.04 | 0.01 | 0.00 | 0.00 | 0.00 | 0.00 | 0.00 | 0.00 |
| | $^{pPgk1}$Foxg1-OE | 0.34 | 0.37 | 0.21 | 0.07 | 0.01 | 0.00 | 0.00 | 0.00 | 0.00 | 0.00 |
| | expr decile | 10 | 9 | 8 | 7 | 6 | 5 | 4 | 3 | 2 | 1 |

| NP | $^{pPgk1}$Plap-OE cell n = 898 | | $^{pPgk1}$Foxg1-OE cell n = 1967 | | | | | | | |
|---|---|---|---|---|---|---|---|---|---|---|
| cell fraction | $^{pPgk1}$Plap-OE | 0.00 | 0.00 | 0.02 | 0.07 | 0.13 | 0.16 | 0.17 | 0.20 | 0.15 | 0.08 |
| | $^{pPgk1}$Foxg1-OE | 0.00 | 0.02 | 0.11 | 0.14 | 0.12 | 0.09 | 0.08 | 0.11 | 0.09 | 0.24 |
| | expr decile | 10 | 9 | 8 | 7 | 6 | 5 | 4 | 3 | 2 | 1 |

| N | $^{pPgk1}$Plap-OE cell n = 2034 | | $^{pPgk1}$Foxg1-OE cell n = 3542 | | | | | | | |
|---|---|---|---|---|---|---|---|---|---|---|
| cell fraction | $^{pPgk1}$Plap-OE | 0.05 | 0.10 | 0.14 | 0.13 | 0.10 | 0.09 | 0.08 | 0.07 | 0.09 | 0.13 |
| | $^{pPgk1}$Foxg1-OE | 0.07 | 0.14 | 0.14 | 0.10 | 0.08 | 0.05 | 0.05 | 0.05 | 0.06 | 0.24 |
| | expr decile | 10 | 9 | 8 | 7 | 6 | 5 | 4 | 3 | 2 | 1 |

### *Figure 6*

**A**

| Foxg1@mLINE1.5'UTR.A, IgG/Plap-OE-norm enrichment | aFoxg1 / Plap-OE | IgG / Plap-OE | aFoxg1 / Foxg1-OE | IgG / Foxg1-OE |
|---|---|---|---|---|
| | 1.60 | 1.34 | 1.71 | 0.64 |
| | 1.08 | 1.39 | 0.90 | 0.26 |
| | 0.90 | 0.48 | 1.85 | 0.60 |
| | 2.03 | 0.78 | 1.21 | 1.05 |

| | |
|---|---|
| IgG/Plap-OE, abs enrichment | 0.0023 |

| Foxg1@mLINE1.5'UTR.Gf, IgG/Plap-OE-norm enrichment | aFoxg1 / Plap-OE | IgG / Plap-OE | aFoxg1 / Foxg1-OE | IgG / Foxg1-OE |
|---|---|---|---|---|
| | 1.43 | 1.33 | 1.41 | 0.60 |
| | 0.93 | 1.33 | 1.11 | 0.24 |
| | 1.02 | 0.53 | 2.04 | 0.61 |
| | 1.99 | 0.81 | 1.29 | 1.05 |

| | |
|---|---|
| IgG/Plap-OE, abs enrichment | 0.0024 |

| Foxg1@mLINE1.5'UTR.Tf, IgG/Plap-OE-norm enrichment | aFoxg1 / Plap-OE | IgG / Plap-OE | aFoxg1 / Foxg1-OE | IgG / Foxg1-OE |
|---|---|---|---|---|
| | 1.78 | 1.35 | 1.74 | 0.61 |
| | 1.12 | 1.36 | 1.26 | 0.28 |
| | 0.96 | 0.56 | 1.77 | 0.61 |
| | 2.09 | 0.73 | 1.24 | 0.97 |

| | |
|---|---|
| IgG/Plap-OE, abs enrichment | 0.0020 |

| Foxg1@mLINE1.orf2, IgG/Plap-OE-norm enrichment | aFoxg1 / Plap-OE | IgG / Plap-OE | aFoxg1 / Foxg1-OE | IgG / Foxg1-OE |
|---|---|---|---|---|
| | | 1.05 | 2.02 | 0.74 |
| | 0.81 | 2.12 | 1.36 | 0.50 |
| | 1.25 | 0.34 | 2.70 | 1.02 |
| | 1.04 | 0.49 | 2.97 | 2.07 |

| | |
|---|---|
| IgG/Plap-OE, abs enrichment | 0.0012 |

| Foxg1@mLINE1.3'UTR, IgG/Plap-OE-norm enrichment | aFoxg1 / Plap-OE | IgG / Plap-OE | aFoxg1 / Foxg1-OE | IgG / Foxg1-OE |
|---|---|---|---|---|
| | 1.16 | 0.98 | 1.43 | 0.65 |
| | 0.76 | 1.61 | 1.15 | 0.18 |
| | 0.97 | 0.66 | 1.75 | 0.66 |
| | 3.46 | 0.74 | 1.15 | 1.03 |

| | |
|---|---|
| IgG/Plap-OE, abs enrichment | 0.0032 |

**B**

| Foxg1@mLINE1.5'UTR.A, IgG/Plap-OE-norm enrichment | aFoxg1 / Plap-OE | IgG / Plap-OE | aFoxg1 / Foxg1-OE | IgG / Foxg1-OE |
|---|---|---|---|---|
| | 3.49 | 1.81 | | 1.13 |
| | 1.62 | 0.34 | 3.74 | 1.62 |
| | 1.97 | 0.45 | 7.47 | 4.03 |
| | 2.72 | 1.41 | 3.54 | 0.60 |
| | 3.05 | 0.99 | 3.88 | 2.83 |

| | |
|---|---|
| IgG/Plap-OE, abs enrichment | 0.0044 |

| Foxg1@mLINE1.5'UTR.Gf, IgG/Plap-OE-norm enrichment | aFoxg1 / Plap-OE | IgG / Plap-OE | aFoxg1 / Foxg1-OE | IgG / Foxg1-OE |
|---|---|---|---|---|
| | 2.63 | | | |
| | 1.01 | 1.24 | 2.45 | 0.82 |
| | 1.41 | 0.63 | 6.76 | 0.86 |
| | 3.27 | 1.27 | 3.02 | 0.64 |
| | 2.49 | 0.86 | 3.13 | 2.03 |

| | |
|---|---|
| IgG/Plap-OE, abs enrichment | 0.0130 |

| Foxg1@mLINE1.5'UTR.Tf, IgG/Plap-OE-norm enrichment | aFoxg1 / Plap-OE | IgG / Plap-OE | aFoxg1 / Foxg1-OE | IgG / Foxg1-OE |
|---|---|---|---|---|
| | 3.34 | 1.60 | | 1.33 |
| | 1.44 | 0.39 | 4.44 | 1.58 |
| | 1.57 | 0.45 | 9.54 | 4.82 |
| | 3.84 | 1.64 | 3.21 | 0.62 |
| | 2.98 | 0.92 | 3.82 | 3.02 |

| | |
|---|---|
| IgG/Plap-OE, abs enrichment | 0.0040 |

| Foxg1@mLINE1.orf2, IgG/Plap-OE-norm enrichment | aFoxg1 / Plap-OE | IgG / Plap-OE | aFoxg1 / Foxg1-OE | IgG / Foxg1-OE |
|---|---|---|---|---|
| | 4.45 | 1.82 | | 1.38 |
| | 0.95 | 0.19 | 3.68 | 1.50 |
| | 1.59 | 0.47 | 13.86 | 6.57 |
| | 4.07 | 1.57 | 5.20 | 0.62 |
| | 3.48 | 0.95 | 5.02 | 3.61 |

| | |
|---|---|
| IgG/Plap-OE, abs enrichment | 0.0011 |

| Foxg1@mLINE1.3'UTR, IgG/Plap-OE-norm enrichment | aFoxg1 / Plap-OE | IgG / Plap-OE | aFoxg1 / Foxg1-OE | IgG / Foxg1-OE |
|---|---|---|---|---|
| | 2.54 | 1.48 | | 0.92 |
| | 1.37 | 0.48 | 1.41 | 0.98 |
| | 1.68 | 0.67 | 5.25 | 3.27 |
| | 2.48 | 1.35 | 3.12 | 0.80 |
| | 2.56 | 1.02 | 2.76 | 2.22 |

| | |
|---|---|
| IgG/Plap-OE, abs enrichment | 0.0060 |

*Figure 8*

**C**

### H3K4me3

**mLINE1.5'UTR.A**

| H3K4me3@mLINE1.5'UTR.A, IgG/Plap-OE-norm enrichment | aH3K4me3 / Plap-OE | IgG / Plap-OE | aH3K4me3 / Foxg1-OE | IgG / Foxg1-OE |
|---|---|---|---|---|
| | 71.5 | 1.5 | | |
| | 82.6 | 0.5 | 66.6 | 1.1 |
| | 76.6 | 1.2 | 59.7 | 0.9 |
| | 76.3 | 0.8 | 71.7 | 0.9 |
| IgG/Plap-OE, abs enrichment | 0.001149219 | | | |

**mLINE1.5'UTR.Gf**

| H3K4me3@mLINE1.5'UTR.Gf, IgG/Plap-OE-norm enrichment | aH3K4me3 / Plap-OE | IgG / Plap-OE | aH3K4me3 / Foxg1-OE | IgG / Foxg1-OE |
|---|---|---|---|---|
| | 103.0 | 1.6 | | |
| | 148.3 | 0.5 | 119.5 | 1.0 |
| | 125.4 | 1.3 | 89.4 | 1.1 |
| | 119.5 | 0.7 | 106.6 | 1.0 |
| IgG/Plap-OE, abs enrichment | 0.000689949 | | | |

**mLINE1.5'UTR.Tf**

| H3K4me3@mLINE1.5'UTR.Tf, IgG/Plap-OE-norm enrichment | aH3K4me3 / Plap-OE | IgG / Plap-OE | aH3K4me3 / Foxg1-OE | IgG / Foxg1-OE |
|---|---|---|---|---|
| | 116.1 | 1.6 | | 1.1 |
| | 146.4 | 0.5 | 117.4 | 1.2 |
| | 118.8 | 1.2 | 81.2 | 0.7 |
| | 103.3 | 0.7 | 113.1 | 1.0 |
| IgG/Plap-OE, abs enrichment | 0.000750664 | | | |

**mLINE1.orf2**

| H3K4me3@mLINE1.orf2, IgG/Plap-OE-norm enrichment | aH3K4me3 / Plap-OE | IgG / Plap-OE | aH3K4me3 / Foxg1-OE | IgG / Foxg1-OE |
|---|---|---|---|---|
| | | 1.4 | | |
| | 163.2 | 0.5 | 124.5 | 0.8 |
| | 193.4 | 1.3 | 108.8 | 1.0 |
| | 152.2 | 0.8 | 163.1 | 0.9 |
| IgG/Plap-OE, abs enrichment | 0.000466815 | | | |

**mLINE1.3'UTR**

| H3K4me3@mLINE1.3'UTR, IgG/Plap-OE-norm enrichment | aH3K4me3 / Plap-OE | IgG / Plap-OE | aH3K4me3 / Foxg1-OE | IgG / Foxg1-OE |
|---|---|---|---|---|
| | 75.8 | 1.6 | | 1.0 |
| | 105.9 | 0.4 | 82.5 | 1.0 |
| | 85.2 | 1.2 | 61.3 | 0.6 |
| | 83.4 | 0.7 | 47.8 | 0.5 |
| IgG/Plap-OE, abs enrichment | 0.000820073 | | | |

### H3K9me3

**mLINE1.5'UTR.A**

| H3K9me3@mLINE1.5'UTR.A, IgG/Plap-OE-norm enrichment | aH3K9me3 / Plap-OE | IgG / Plap-OE | aH3K9me3 / Foxg1-OE | IgG / Foxg1-OE |
|---|---|---|---|---|
| | 99.9 | | 112.6 | 0.7 |
| | 91.6 | 1.3 | 91.1 | 0.6 |
| | 82.0 | 0.7 | 86.3 | 0.5 |
| | 65.3 | 1.0 | 85.2 | 1.1 |
| | 73.7 | 1.0 | 155.1 | 1.6 |
| IgG/Plap-OE, abs enrichment | 0.002083091 | | | |

**mLINE1.5'UTR.Gf**

| H3K9me3@mLINE1.5'UTR.Gf, IgG/Plap-OE-norm enrichment | aH3K9me3 / Plap-OE | IgG / Plap-OE | aH3K9me3 / Foxg1-OE | IgG / Foxg1-OE |
|---|---|---|---|---|
| | 79.4 | | 94.0 | 0.7 |
| | 81.0 | 1.2 | 83.6 | 0.7 |
| | 79.5 | 0.8 | 88.7 | 0.7 |
| | 83.6 | 1.0 | 104.0 | 1.2 |
| | 78.6 | 1.0 | 88.6 | 1.6 |
| IgG/Plap-OE, abs enrichment | 0.002339303 | | | |

**mLINE1.5'UTR.Tf**

| H3K9me3@mLINE1.5'UTR.Tf, IgG/Plap-OE-norm enrichment | aH3K9me3 / Plap-OE | IgG / Plap-OE | aH3K9me3 / Foxg1-OE | IgG / Foxg1-OE |
|---|---|---|---|---|
| | | | | 0.6 |
| | 100.1 | | 118.0 | 0.6 |
| | 89.8 | 1.1 | 91.4 | 0.6 |
| | 81.6 | 0.9 | 91.4 | 1.0 |
| | 82.6 | 1.0 | 110.7 | 1.3 |
| IgG/Plap-OE, abs enrichment | 0.002191176 | | | |

**mLINE1.orf2**

| H3K9me3@mLINE1.orf2, IgG/Plap-OE-norm enrichment | aH3K9me3 / Plap-OE | IgG / Plap-OE | aH3K9me3 / Foxg1-OE | IgG / Foxg1-OE |
|---|---|---|---|---|
| | 167.7 | | 210.7 | 0.6 |
| | 150.6 | 1.5 | 216.4 | 0.6 |
| | 165.6 | 0.6 | 168.3 | 0.5 |
| | 119.5 | 0.9 | 156.2 | 1.2 |
| | 71.6 | 0.9 | 180.7 | 1.7 |
| IgG/Plap-OE, abs enrichment | 0.000911135 | | | |

**mLINE1.3'UTR**

| H3K9me3@mLINE1.3'UTR, IgG/Plap-OE-norm enrichment | aH3K9me3 / Plap-OE | IgG / Plap-OE | aH3K9me3 / Foxg1-OE | IgG / Foxg1-OE |
|---|---|---|---|---|
| | 63.2 | | 70.6 | 0.7 |
| | 61.4 | 1.3 | 58.6 | 0.6 |
| | 58.8 | 0.7 | 59.9 | 0.7 |
| | 45.1 | 1.0 | 69.2 | 1.2 |
| | 54.1 | 1.0 | 78.2 | 1.7 |
| IgG/Plap-OE, abs enrichment | 0.002530524 | | | |

##### H3K27aac

**mLINE1.5'UTR.A**

| H3K27ac@mLINE1.5'UTR.A, IgG/Plap-OE-norm enrichment | aH3K27ac / Plap-OE | IgG / Plap-OE | aH3K27ac / Foxg1-OE | IgG / Foxg1-OE |
|---|---|---|---|---|
| | 530.7 | | 491.3 | 1.8 |
| | 489.0 | 0.9 | 554.3 | 0.0 |
| | 458.0 | 1.2 | 607.0 | 2.2 |
| | 484.4 | 0.7 | 455.2 | 3.3 |
| | 524.5 | 1.3 | 605.2 | 2.7 |
| IgG/Plap-OE, abs enrichment | 0.001826347 | | | |

**mLINE1.5'UTR.Gf**

| H3K27ac@mLINE1.5'UTR.Gf, IgG/Plap-OE-norm enrichment | aH3K27ac / Plap-OE | IgG / Plap-OE | aH3K27ac / Foxg1-OE | IgG / Foxg1-OE |
|---|---|---|---|---|
| | 1159.5 | | 1187.2 | 2.3 |
| | 1092.1 | 0.7 | 1110.7 | 0.0 |
| | 1272.5 | 1.1 | 1261.2 | 2.9 |
| | 1142.8 | 0.6 | 785.1 | 3.4 |
| | 969.9 | 1.5 | 1500.0 | 2.0 |
| IgG/Plap-OE, abs enrichment | 0.001087686 | | | |

**mLINE1.5'UTR.Tf**

| H3K27ac@mLINE1.5'UTR.Tf, IgG/Plap-OE-norm enrichment | aH3K27ac / Plap-OE | IgG / Plap-OE | aH3K27ac / Foxg1-OE | IgG / Foxg1-OE |
|---|---|---|---|---|
| | 1104.1 | | | 2.0 |
| | 1025.9 | 0.9 | 1221.4 | 0.0 |
| | 1219.0 | 1.1 | 1131.1 | 3.1 |
| | 1074.1 | 0.7 | 1355.9 | 3.3 |
| | 962.4 | 1.3 | 799.9 | 2.1 |
| IgG/Plap-OE, abs enrichment | 0.000980244 | | | |

**mLINE1.orf2**

| H3K27ac@mLINE1.orf2, IgG/Plap-OE-norm enrichment | aH3K27ac / Plap-OE | IgG / Plap-OE | aH3K27ac / Foxg1-OE | IgG / Foxg1-OE |
|---|---|---|---|---|
| | 1028.8 | | | 3.1 |
| | 848.1 | 0.9 | 955.9 | 0.0 |
| | 933.1 | 0.9 | 922.0 | 3.9 |
| | 893.7 | 0.7 | 608.1 | 2.6 |
| | 814.7 | 1.5 | 1121.4 | 2.0 |
| IgG/Plap-OE, abs enrichment | 0.001110746 | | | |

**mLINE1.3'UTR**

| H3K27ac@mLINE1.3'UTR, IgG/Plap-OE-norm enrichment | aH3K27ac / Plap-OE | IgG / Plap-OE | aH3K27ac / Foxg1-OE | IgG / Foxg1-OE |
|---|---|---|---|---|
| | 1690.3 | 0.0 | 1402.9 | 2.2 |
| | 1059.0 | 0.8 | 1162.5 | 0.0 |
| | 1190.2 | 1.1 | 1442.5 | 3.5 |
| | 1530.8 | 0.6 | 926.3 | 3.4 |
| | 1107.3 | 1.6 | 1131.6 | 1.4 |
| IgG/Plap-OE, abs enrichment | 0.001167836 | | | |

##### MeCP2

**mLINE1.5'UTR.A**

| MeCP2@mLINE1.5'UTR.A, IgG/Plap-OE-norm enrichment | aMeCP2 / Plap-OE | IgG / Plap-OE | aMeCP2 / Foxg1-OE | IgG / Foxg1-OE |
|---|---|---|---|---|
| | 2.1 | 1.1 | 1.6 | 1.3 |
| | 1.5 | 0.9 | 2.0 | 1.8 |
| | 1.1 | 1.0 | 1.6 | 1.4 |
| | 1.6 | 1.0 | 3.5 | 2.9 |
| IgG/Plap-OE, abs enrichment | 0.000934096 | | | |

**mLINE1.5'UTR.Gf**

| MeCP2@mLINE1.5'UTR.Gf, IgG/Plap-OE-norm enrichment | aMeCP2 / Plap-OE | IgG / Plap-OE | aMeCP2 / Foxg1-OE | IgG / Foxg1-OE |
|---|---|---|---|---|
| | 2.5 | | | |
| | 1.7 | | 1.4 | 1.3 |
| | 1.0 | 1.1 | 2.1 | 1.6 |
| | 1.2 | 0.8 | 1.6 | 1.3 |
| | 1.7 | 1.1 | 3.0 | 2.4 |
| IgG/Plap-OE, abs enrichment | 0.000777261 | | | |

**mLINE1.5'UTR.Tf**

| MeCP2@mLINE1.5'UTR.Tf, IgG/Plap-OE-norm enrichment | aMeCP2 / Plap-OE | IgG / Plap-OE | aMeCP2 / Foxg1-OE | IgG / Foxg1-OE |
|---|---|---|---|---|
| | 2.4 | | | |
| | 1.7 | 1.0 | 1.9 | 1.3 |
| | 1.1 | 1.0 | 1.6 | 2.2 |
| | 1.5 | 1.0 | 3.1 | 1.8 |
| | 2.1 | 0.9 | 5.0 | 3.3 |
| IgG/Plap-OE, abs enrichment | 0.001044369 | | | |

**mLINE1.orf2**

| MeCP2@mLINE1.orf2, IgG/Plap-OE-norm enrichment | aMeCP2 / Plap-OE | IgG / Plap-OE | aMeCP2 / Foxg1-OE | IgG / Foxg1-OE |
|---|---|---|---|---|
| | 1.4 | | | |
| | 1.0 | 0.9 | 1.3 | 0.9 |
| | 0.8 | 1.1 | 1.3 | 1.2 |
| | 1.0 | 1.1 | 1.8 | 1.8 |
| | 1.3 | 0.9 | 2.3 | 1.7 |
| IgG/Plap-OE, abs enrichment | 0.001918216 | | | |

**mLINE1.3'UTR**

| MeCP2@mLINE1.3'UTR, IgG/Plap-OE-norm enrichment | aMeCP2 / Plap-OE | IgG / Plap-OE | aMeCP2 / Foxg1-OE | IgG / Foxg1-OE |
|---|---|---|---|---|
| | 2.1 | | | |
| | 1.6 | 0.8 | 1.8 | 1.4 |
| | 1.1 | 1.0 | 1.7 | 2.1 |
| | 1.4 | 1.3 | 3.3 | 1.9 |
| | 1.9 | 0.9 | 3.2 | 2.8 |
| IgG/Plap-OE, abs enrichment | 0.000483773 | | | |

*Figure 9*

| "quadruplet", Plap-OE double-normalized *mLINE1-mRNA* | Foxg1 (W308X) | PLAP | Foxg1 (W308X) | PLAP | Foxg1 (W308X) | PLAP | Foxg1 (W308X) | PLAP |
|---|---|---|---|---|---|---|---|---|
| | *mLINE1-5'UTR.A* | | *mLINE1-5'UTR.Gf* | | *mLINE1-5'UTR.Tf* | | *mLINE1 Orf2* | |
| | 0.77 | 0.99 | 1.00 | 1.04 | 0.81 | 1.03 | 0.75 | 1.08 |
| | 0.91 | 1.09 | 1.01 | 1.13 | 1.05 | 1.06 | 0.78 | 1.00 |
| | 0.82 | 1.06 | 0.87 | 0.93 | 0.85 | 1.02 | 0.69 | 0.91 |
| | 0.80 | 0.99 | 0.79 | 0.99 | 0.91 | 0.96 | 0.81 | 1.00 |
| | 0.97 | 0.88 | 1.14 | 0.91 | 0.97 | 0.93 | 1.01 | 1.01 |

*Figure 10*

B1

| [Gfap-S100β, Foxg1] ctrl-double-norm *mLINE1-3'UTR DNA* | mLine1.3'UTR | | |
|---|---|---|---|
| | Prot I | Prot II | Prot III |
| | | | 1.31 |
| | | 1.12 | 1.36 |
| | 0.89 | 1.13 | 1.24 |
| | 1.11 | 1.05 | 1.44 |
| | 1.19 | 1.24 | 1.25 |
| | 1.12 | 0.82 | 1.51 |
| | 0.69 | 0.79 | 1.35 |

B2

| [Gfap-S100β, Foxg1] ctrl-double-norm *mLINE1-3'UTR DNA* | mLine1.3'UTR | | |
|---|---|---|---|
| | E15.5 | E20.5 Lam- | E20.5 Lam+ |
| | 0.99 | 1.00 | 1.00 |
| | 1.15 | 1.02 | 0.88 |
| | 0.85 | 1.66 | 0.90 |
| | 1.05 | 1.11 | 1.15 |
| | 1.00 | 1.14 | 0.91 |
| | 1.03 | 1.26 | 1.00 |
| | 0.97 | 1.05 | 0.85 |
| | 0.97 | 2.25 | 0.93 |

B3

| [Gfap-S100β, Foxg1] ctrl-double-norm *mLINE1-3'UTR DNA* | mLine1.3'UTR | |
|---|---|---|
| | E15.5 | E20.5 |
| | 1.09 | |
| | 1.10 | 1.06 |
| | 0.82 | 1.31 |
| | 0.94 | 0.73 |
| | 1.02 | 0.96 |
| | 1.07 | 1.25 |
| | 1.08 | 1.27 |
| | 0.89 | 1.39 |

B4

| [Gfap-S100β, Foxg1] ctrl-double-norm *mLINE1-3'UTR DNA* | mLine1.3'UTR | |
|---|---|---|
| | E15.5 | E20.5 |
| | 0.93 | 0.97 |
| | 0.92 | 1.09 |
| | 0.96 | 0.81 |
| | 1.00 | 1.27 |
| | 0.99 | 1.48 |
| | 1.14 | 1.15 |
| | 1.05 | 1.06 |

B5

| [Gfap-S100β, Foxg1] ctrl-double-norm *mLINE1-3'UTR DNA* | mLine1.3'UTR | |
|---|---|---|
| | E15.5 | E20.5 |
| | 0.97 | |
| | 0.97 | 0.99 |
| | 0.83 | 1.27 |
| | 1.04 | 0.73 |
| | 1.06 | 1.04 |
| | 1.00 | 1.18 |
| | 0.95 | 1.11 |
| | 1.17 | 1.12 |

*Figure 11*

*Neocortex*

| [Gfap-S100β, Foxg1] ctrl-double-norm *mLINE1-3'UTR DNA* | mLine1.3'UTR | |
|---|---|---|
| | *Foxg1* $^{+/+}$ E14 | *Foxg1* $^{+/+}$ P0 |
| | 0.72 | |
| | 1.04 | 1.30 |
| | 0.79 | 1.01 |
| | 0.88 | 1.24 |
| | 1.11 | 1.26 |
| | 1.13 | 1.41 |
| | 1.11 | 1.17 |
| | 1.22 | 1.27 |

*Mesencephalon*

| [Gfap-S100β, Foxg1] ctrl-double-norm *mLINE1-3'UTR DNA* | mLine1.3'UTR | |
|---|---|---|
| | *Foxg1* $^{+/+}$ E14 | *Foxg1* $^{+/+}$ P0 |
| | | 1.25 |
| | | 1.45 |
| | | 1.25 |
| | 0.60 | 1.50 |
| | 1.09 | 1.91 |
| | 1.21 | 1.86 |
| | 0.88 | 1.72 |
| | 1.08 | 1.56 |
| | 1.14 | 1.38 |

*Figure 12*

| [Gfap-Nfia] ctrl-double-norm *mLINE1-3'UTR DNA* | mLine1.3'UTR | |
|---|---|---|
| | *Foxg1* $^{+/+}$ P0 | *Foxg1* $^{+/-}$ P0 |
| | 0.96 | |
| | 0.96 | |
| | 0.99 | 0.92 |
| | 0.98 | 0.91 |
| | 1.05 | 0.93 |
| | 1.06 | 0.94 |
| | 0.98 | 0.89 |
| | 1.03 | 0.96 |

*Figure 13*

**B1**

| Gfap- ctrl-double-norm *mLINE1-3'UTR DNA* | | |
|---|---|---|
| | 0.97 | 0.80 |
| | 0.91 | 0.92 |
| | 1.00 | 0.92 |
| | 0.93 | 0.82 |
| | 1.08 | 0.92 |
| | 0.99 | 0.83 |
| | 1.07 | 0.76 |
| | 1.05 | 0.82 |
| | Ctrl | *Foxg1* - LOF |
| | *mLINE1* .3'UTR | |

**B2**

| Gfap- ctrl-double-norm *mLINE1-3'UTR DNA* | | |
|---|---|---|
| | 0.96 | 0.71 |
| | 0.83 | 0.85 |
| | 0.93 | 0.79 |
| | 0.87 | 0.63 |
| | 1.12 | 0.81 |
| | 1.11 | 0.72 |
| | 1.09 | 0.62 |
| | 1.09 | 0.69 |
| | Ctrl | *Foxg1* - LOF |
| | *mLINE1* .3'UTR | |

**B3**

| Gfap- ctrl-double-norm *mLINE1-3'UTR DNA* | | |
|---|---|---|
| | 1.00 | 0.95 |
| | 1.22 | 0.83 |
| | 1.04 | 1.08 |
| | 1.10 | 1.04 |
| | 1.22 | 0.66 |
| | 0.95 | 0.96 |
| | 0.44 | 1.05 |
| | 1.04 | 0.96 |
| | 0.81 | 1.06 |
| | 1.20 | 1.39 |
| | Ctrl | *Foxg1* - OE |
| | *mLINE1* .3'UTR | |

**B4**

| Gfap- ctrl-double-norm *mLINE1-3'UTR DNA* | | |
|---|---|---|
| | 0.89 | 2.43 |
| | 1.17 | 1.44 |
| | 1.26 | 1.44 |
| | 0.64 | 0.89 |
| | 0.82 | 1.40 |
| | 1.18 | 1.33 |
| | 1.15 | 1.24 |
| | 0.62 | 1.52 |
| | 1.29 | 0.65 |
| | Ctrl | *Foxg1* - OE |
| | *mLINE1* .3'UTR | |

*Figure 14*

| *Gfap*-, "prot III"-double-norm DNA | | | | | | |
|---|---|---|---|---|---|---|
| | 1.20 | | | | | |
| | 1.07 | | | | | |
| | 0.97 | | | | | |
| | 1.02 | | | | | |
| | 0.95 | | | | | |
| | 0.88 | | | | | |
| | 0.91 | | | | | |
| | 0.97 | | | | | |
| | 0.97 | | | | | |
| | 0.97 | | | | | |
| | 1.16 | | | | | |
| | 1.15 | | | | | |
| | 1.30 | | | | | |
| | 0.66 | | | | | |
| | 0.82 | | 1.05 | 0.65 | | |
| | 1.06 | | 1.27 | 0.70 | | |
| | 0.83 | 0.64 | 1.06 | 0.88 | | |
| | 1.12 | 0.74 | 1.02 | 0.98 | 0.83 | |
| | 1.16 | 1.15 | 1.20 | 0.85 | 0.94 | 0.89 |
| | 0.94 | 0.90 | 1.27 | 0.78 | 0.87 | 0.88 |
| | 0.95 | 1.03 | 1.20 | 0.74 | 0.99 | 0.89 |
| | 1.00 | 0.96 | 1.13 | 0.67 | 0.83 | 0.94 |
| | 0.93 | 0.98 | 1.18 | 0.53 | 0.92 | 0.97 |
| | 1.01 | 0.97 | 1.31 | 0.58 | 0.73 | 0.81 |
| *Foxg1* | wt | wt | wt | lof | lof | lof |
| *Mov10* | wt | lof | wt | wt | lof | wt |
| *Ddx39a* | wt | wt | lof | wt | wt | lof |
| | *mLine1.* 3'UTR | | | | | |

*Figure 15*

(1)

| Foxg1@mLINE1.5'UTR.A, IgG/Plap-OE-norm enrichment | | |
|---|---|---|
| | 2.68 | |
| | 2.73 | |
| | 3.54 | 0.98 |
| | 2.95 | 0.82 |
| | 3.63 | 1.20 |
| | aFoxg1 / Plap-OE | IgG / Plap-OE |

(2)

| Foxg1@mLINE1.5'UTR.Gf, IgG/Plap-OE-norm enrichment | | |
|---|---|---|
| | 1.49 | |
| | 1.17 | |
| | 2.26 | 0.55 |
| | 1.61 | 1.05 |
| | 2.73 | 1.40 |
| | aFoxg1 / Plap-OE | IgG / Plap-OE |

(3)

| Foxg1@mLINE1.5'UTR.Tf, IgG/Plap-OE-norm enrichment | | |
|---|---|---|
| | 5.34 | |
| | 2.80 | |
| | 3.50 | 0.57 |
| | 3.69 | 0.56 |
| | 5.32 | 1.87 |
| | aFoxg1 / Plap-OE | IgG / Plap-OE |

(4)

| Foxg1@mLINE1.Orf2, IgG/Plap-OE-norm enrichment | | |
|---|---|---|
| | 2.23 | |
| | 0.74 | |
| | 1.06 | 0.93 |
| | 1.95 | 1.13 |
| | 3.34 | 0.95 |
| | aFoxg1 / Plap-OE | IgG / Plap-OE |

(5)

| Foxg1@mLINE1.3'UTR., IgG/Plap-OE-norm enrichment | | |
|---|---|---|
| | 3.81 | |
| | 2.35 | |
| | 3.58 | 0.84 |
| | 3.91 | 1.24 |
| | 5.78 | 0.92 |
| | aFoxg1 / Plap-OE | IgG / Plap-OE |

#### *Figure S1*

**A**

| Gapdh-, ctrl-double-norm *mLINE1*-mRNA | mLine1.A ctr | mLine1.A Foxg1-LOF | mLine1.Gf ctr | mLine1.Gf Foxg1-LOF | mLine1.Tf ctr | mLine1.Tf Foxg1-LOF |
|---|---|---|---|---|---|---|
| | | 1.09 | | 1.14 | | 1.03 |
| | 0.76 | 1.23 | 1.11 | 1.30 | 0.89 | 1.26 |
| | 0.92 | 1.32 | 0.88 | 0.94 | 1.04 | 0.98 |
| | 0.78 | 1.28 | 0.87 | 1.12 | 0.83 | 1.12 |
| | 0.86 | 0.66 | 0.76 | 1.39 | 0.97 | 1.04 |
| | 1.55 | 1.18 | 1.33 | 0.93 | 1.20 | 1.25 |
| | 0.98 | 1.48 | 0.94 | 1.59 | 1.06 | 1.52 |
| | 0.82 | 0.87 | 0.76 | 0.85 | 0.80 | 0.93 |
| | 1.15 | 2.09 | 1.10 | 2.11 | 1.05 | 1.77 |
| | 1.11 | 1.13 | 1.07 | 1.16 | 1.07 | 1.26 |
| | 1.07 | 0.99 | 1.17 | 0.89 | 1.10 | 1.03 |

| ["quadruplet"-, ctrl-double-norm *mLINE1*- mRNA] | mLine1.A ctr | mLine1.A Foxg1-LOF | mLine1.Gf ctr | mLine1.Gf Foxg1-LOF | mLine1.Tf ctr | mLine1.Tf Foxg1-LOF |
|---|---|---|---|---|---|---|
| | | 1.18 | | 1.22 | | 1.11 |
| | 0.77 | 1.35 | 1.11 | 1.41 | 0.89 | 1.37 |
| | 0.89 | 1.43 | 0.84 | 1.00 | 0.99 | 1.05 |
| | 0.97 | 1.58 | 1.07 | 1.37 | 1.03 | 1.38 |
| | 0.83 | 0.86 | 0.73 | 1.79 | 0.93 | 1.34 |
| | 1.31 | 0.77 | 1.12 | 0.60 | 1.01 | 0.81 |
| | 0.83 | 1.09 | 0.80 | 1.17 | 0.90 | 1.12 |
| | 0.82 | 0.82 | 0.76 | 0.80 | 0.80 | 0.88 |
| | 1.02 | 1.31 | 0.97 | 1.31 | 0.93 | 1.11 |
| | 1.32 | 1.09 | 1.27 | 1.11 | 1.26 | 1.20 |
| | 1.23 | 0.93 | 1.34 | 0.84 | 1.26 | 0.96 |

**B**

| Gapdh-, ctrl-double-norm *mLINE1*-mRNA | mLine1.orf2 ctr | mLine1.orf2 Foxg1-OE | mLine1.A ctr | mLine1.A Foxg1-OE | mLine1.Gf ctr | mLine1.Gf Foxg1-OE | mLine1.Tf ctr | mLine1.Tf Foxg1-OE |
|---|---|---|---|---|---|---|---|---|
| | 0.94 | 0.85 | 0.84 | 0.35 | 0.96 | 0.31 | 0.86 | 0.43 |
| | 0.58 | 0.60 | 0.80 | 0.28 | 0.77 | 0.25 | 0.93 | 0.34 |
| | 1.64 | 0.33 | 1.30 | 0.23 | 1.37 | 0.22 | 1.43 | 0.24 |
| | 1.12 | 0.68 | 1.03 | 0.56 | 1.14 | 0.51 | 1.21 | 0.76 |
| | 1.15 | 0.41 | 1.06 | 0.47 | 0.88 | 0.26 | 0.86 | 0.30 |
| | 0.57 | 0.66 | 0.98 | 0.41 | 0.88 | 0.40 | 0.72 | 0.39 |

| ["quadruplet"-, *Plap*-OE-double norm mLINE1- mRNA] | mLine1.orf2 ctr | mLine1.orf2 Foxg1-OE | mLine1.A ctr | mLine1.A Foxg1-OE | mLine1.Gf ctr | mLine1.Gf Foxg1-OE | mLine1.Tf ctr | mLine1.Tf Foxg1-OE |
|---|---|---|---|---|---|---|---|---|
| | 0.95 | 0.74 | 0.87 | 0.31 | 0.99 | 0.28 | 0.89 | 0.38 |
| | 0.48 | 0.72 | 0.67 | 0.35 | 0.65 | 0.30 | 0.78 | 0.41 |
| | 1.96 | 0.35 | 1.60 | 0.26 | 1.67 | 0.24 | 1.75 | 0.26 |
| | 1.04 | 0.57 | 0.98 | 0.48 | 1.08 | 0.44 | 1.14 | 0.65 |
| | 1.09 | 0.46 | 1.03 | 0.54 | 0.85 | 0.29 | 0.83 | 0.34 |
| | 0.48 | 0.58 | 0.85 | 0.37 | 0.76 | 0.36 | 0.62 | 0.35 |

#### *Figure S2*

*high Pk*

(1)

| *Gfap* -,ctrl-double-norm *Mecp2 DNA* | E15.5 males | E15,5 females |
|---|---|---|
| | 0.97 | |
| | 1.28 | 1.72 |
| | 0.75 | 1.36 |

*very high Pk*

(2)

| *Gfap*, - ctrl-double-norm *Mecp2 DNA* | E15.5 males | E15,5 females |
|---|---|---|
| | 1.02 | |
| | 1.04 | 1.80 |
| | 0.94 | 2.69 |

*very high Pk*

(3)

| *Gfap*, - ctrl-double-norm *Cdkl5 DNA* | E15.5 males | E15,5 females |
|---|---|---|
| | 0.73 | 2.04 |
| | 1.23 | 2.26 |
| | 1.04 | 2.37 |
